# Probing mechanisms of transcription elongation through cell-to-cell variability of RNA polymerase

**DOI:** 10.1101/655712

**Authors:** Md. Zulfikar Ali, Sandeep Choubey, Dipjyoti Das, Robert C. Brewster

## Abstract

The process of transcription initiation and elongation are primary points of control in the regulation of gene expression. While biochemical studies have uncovered the mechanisms involved in controlling transcription at each step, how these mechanisms manifest *in vivo* at the level of individual genes is still unclear. Recent experimental advances have enabled single-cell measurements of RNAP molecules engaged in the process of transcribing a gene of interest. In this manuscript, we use Gillespie simulations to show that measurements of cell-to-cell variability of RNAP numbers and inter-polymerase distances can reveal the prevailing mode of regulation of a given gene. Mechanisms of regulation at each step, from initiation to elongation dynamics, produce qualitatively distinct signatures which can further be used to discern between them. Intriguingly, depending on the initiation kinetics, stochastic elongation can either enhance or suppress cell-to-cell variability at the RNAP level. To demonstrate the value of this framework, we analyze RNAP number distribution data for ribosomal genes in S. cerevisiae from three previously published studies and show that this approach provides crucial mechanistic insights into the transcriptional regulation of these genes.

**Author Summary:** The process of transcription comprises many distinct steps and understanding the regulation of each of these steps provides insight into how the levels of gene expression are controlled in the cell. In this manuscript, we use stochastic simulations to explore how regulation at the level of elongation and initiation together influences the distribution of actively transcribing RNAP on a gene. We find that each of these steps of regulation leaves a distinct imprint on the gene-to-gene variability of number of actively transcribing RNAP on the gene and their inter-RNAP distances. Using these results, we analyze recent experimental data of transcribing RNAP distributions and find that the perturbations in these studies primarily impact the transcription initiation dynamics of the genes.

## Introduction

Transcription broadly consists of three steps: Initiation, elongation, and termination [1,2]. It has become evident that each of these steps can play an important role in the regulation of gene expression to varying degrees [3]. Experimental [4–6] and theoretical [7–9] approaches have typically studied these mechanisms by directly monitoring them individually to gain a mechanistic understanding of each. In this manuscript we aim to demonstrate how single-cell measurements of the number of active, transcribing RNAP can be used to infer the regulatory dynamics of the initiation and elongation process that is controlling the expression of that gene.

Recent experimental advancements have stimulated the analysis of the mechanisms of transcription elongation *in vivo*. Electron micrograph (EM) images [10–13] enable counting nascent RNAs i.e. the number of RNAP molecules engaged in the elongation process of any one gene (shown in Fig. 1B). Similar quantitative information is obtained from Fluorescence *In Situ* Hybridization (FISH) experiments [4,14–16], although distinguishing number of nascent RNAs from the raw signal intensity presents a challenge. Moreover, EM images allow us to extract the inter-RNAP distances along a gene (shown in Fig. 1C). These two kinds of measurements, i.e. the number of transcribing RNAPs and the inter-RNAP distances, obtained from these abovementioned experiments have revealed how different transcription elongation factors such as Spt5 [17] and Nus-factors [18] can affect elongation of ribosomal genes in both *E. coli* and yeast. However, this manuscript lays a foundation for using the variability in these measurements as a way to reveal the specific kinetics of regulation of the measured genes.

**Fig. 1:**
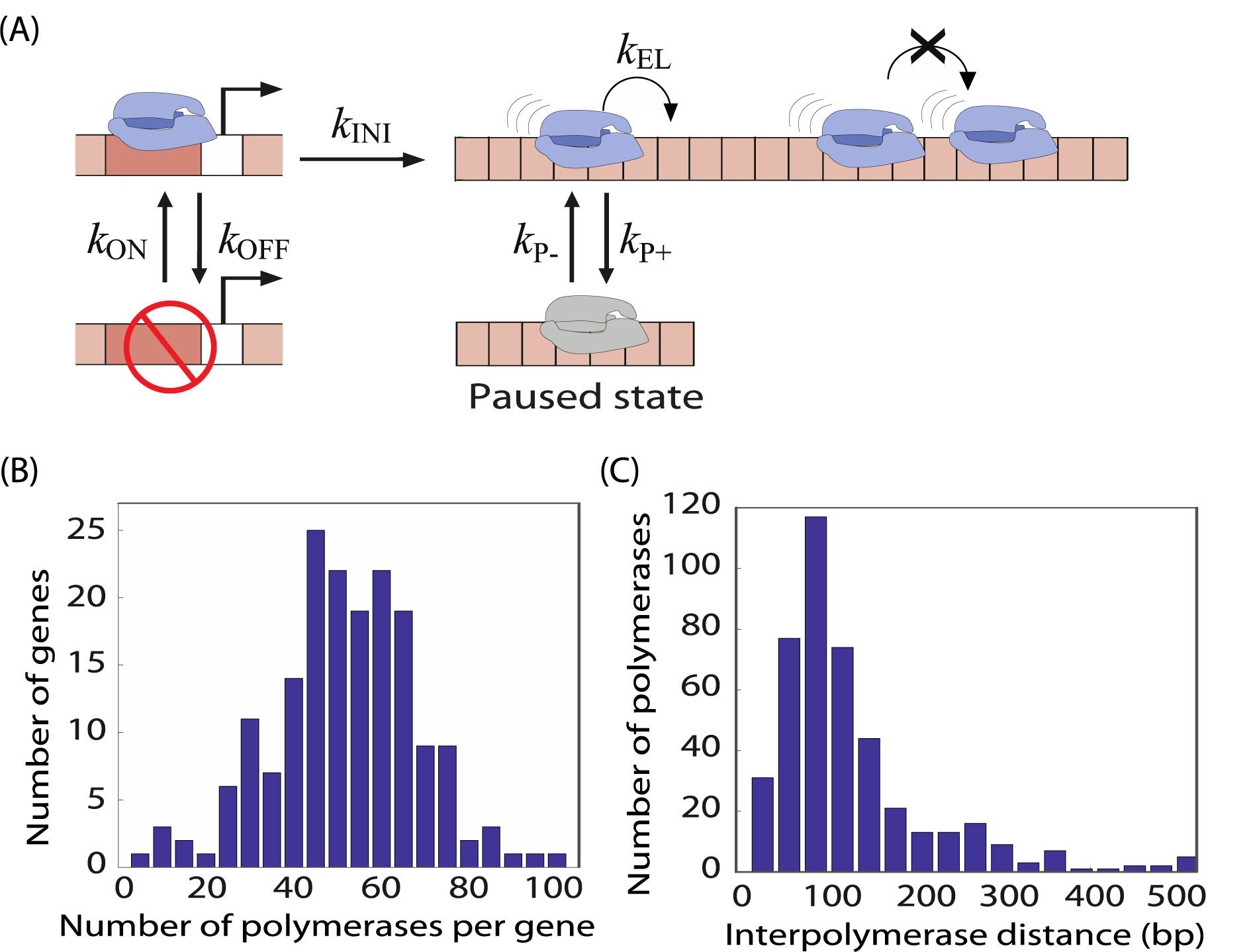
Model of transcription elongation. (A) The promoter for the gene on interest switches between two states: an active and an inactive one. The rate of switching from the active state to the inactive state is *k*_OFF_, and from the inactive to the active state is *k*_ON_. From the active state transcription initiation proceeds at a rate *k*_INI_. Once on the gene, each polymerase molecule can either be in an active or a paused state, independent of the state of other RNAPs. A polymerase molecule in the active state can switch stochastically to the paused state with a rate *k*_P+_, whereas it can again switch back to the active state with rate *k*_P-_. In the active state, the polymerases move from one base pair to the next at a rate *k*_EL_, until they reach the end of the gene and fall off at the same rate. From this model, we compute the mean and the variance of the number of RNA polymerases, in steady state, and the inter-polymerase distances along the gene as a function of the initiation rate. (B) Histogram of the RNAP number distribution is shown for *rrn* gene in yeast (adapted from [17]) obtained from EM images of the DNA extracted from single cells. (C) Histogram of inter-polymerase distances along a gene in *E. coli* (adapted from [18]). EM images also provide inter-polymerase distances along a gene of interest.

There is a history of using kinetic models to describe the dynamic processes of transcriptional regulation [19,20] In particular these models have been used to inform how the mean transcriptional levels and the variability around that mean depends on the dynamics of the processes that regulate transcription. This has culminated in a prevalent “On-off” model [14,19,21–28] of promoter dynamics that considers initiation as being regulated by switching between a transcriptionally active and inactive state, the origin of this switching has been proposed as related to mechanisms such as transcription factor dynamics, chromatin accessibility or condensates of transcriptional machinery. This model has been extended to include the dynamics of the elongation process, to varying degrees; the roles of simple stochastic hopping, pausing and steric hindrance have all been considered and their contribution to phenomena such as traffic jams and transcription rate has been examined. These models of elongation dynamics have been used to predict how mRNA and protein distributions [29,30] depend on elongation dynamics. In light of the data introduced above, the goal of this manuscript is to develop the theoretical predictions that can be used, in conjunction with single-cell transcribing RNAP distributions, to reveal the molecular mechanisms that control transcription of a given gene. Although previous studies have used mRNA and protein distributions to infer these mechanisms, these measurements are conflated with the stochasticity of downstream processes such as translation, post-processing, degradation and partitioning [30–37]. In this work we use direct measurement of nascent mRNA distributions which are not subject to these additional sources of variability.

We find that different mechanisms of elongation lead to distinct signatures at the level of transcribing RNAP numbers and inter-polymerase distances, which can further be used to discriminate between these mechanisms. We also find that for bursty transcription initiation (characterized by periods of high initiation activity followed by extended periods of silence [33]), elongation dynamics enhance cell-to-cell variability (noise) in RNAP numbers for low initiation rates. However, for high initiation rates, elongation dynamics suppresses noise, leading to a masking of the bursty initiation dynamics. To further show the utility of our theoretical framework, we analyze published distributions of RNAP numbers for ribosomal genes in budding yeast (*S. cerevisiae*) and show that in each case the observed effect on the gene-to-gene variability in RNAP numbers, for each perturbation, is consistent with bursty initiation mechanism for those genes.

## Methods

### Model

To demonstrate how the distribution of RNAP numbers and inter-polymerase distances can be used to extract dynamical information about the process of transcription elongation *in vivo*, we examine a simple model of transcription (shown in Fig. 1A) which incorporates both promoter and elongation dynamics. The promoter can transition between two states: an active state (ON), from which transcription initiation can occur, and an inactive state (OFF) from which initiation does not occur through mechanisms such as transcription factor binding [8,10,81–89]. The rate of switching from the active to the inactive state is *k*_OFF_ and inactive to the active state is *k*_ON_, while the rate of initiation from the active state is *k*_INI_. Next, we consider transcription elongation. *In vivo* [6,34–51] and *in vitro* [52–70] experimental studies have reported the asynchronous nature of the motion of RNAP molecules during elongation which includes ubiquitous pausing and stochastic hopping. To incorporate these features of RNAP elongation in the model, we assume that each individual RNAP molecule on the gene transitions between two states: an active state, where it stochastically processes (through single base pair hops) down the gene at a rate *k*_EL_, called the hopping rate, or a paused state, where hopping ceases. Similar models [29,39–41] have been developed to capture the stochasticity of elongation dynamics. An RNAP molecule in the active state can switch stochastically to the paused state with a rate *k*_P+_, whereas it can again switch back to the active state with rate *k*_P-_. For elongation, we explicitly consider two different scenarios. First, we study the limit when the RNAPs do not pause along the gene (*k*_P+_→ 0 in Fig 1A) and elongation is governed by the hopping rate, *k*_EL_. We call this limiting case the ***simple elongation model***. In the other scenario, RNAPs pause along the gene at random sites and elongation is characterized by three processes, namely forward hopping, pausing (at rate *k*_P+_), and unpausing (at rate *k*_P-_) of the RNAPs. This second case is referred to as the ***pausing model***. The polymerases are treated as extended, solid objects of length *L*_P_ base-pairs and so any reaction that would create an overlap is forbidden. We assume that each polymerase can pause at any base along the gene and the rate at which it pauses is independent of the identity of the base. However, inclusion of site-specific pausing [41–43] does not change the conclusions of our work and, as such, are primarily presented in the supplementary materials. Additionally, there are several other mechanisms that can affect the progression of RNAP on a gene, e.g., backtracking of polymerases [74, 75], crowding by nucleosome [44], and DNA supercoiling [45,46]. However, to keep the model simple and to reduce the parameter space, we do not incorporate these additional mechanisms. Model parameters used in the simulation are listed in table 1 and 2 and are taken from previously reported papers. We discuss in detail the parameter selection in method section.

**Table 1:**
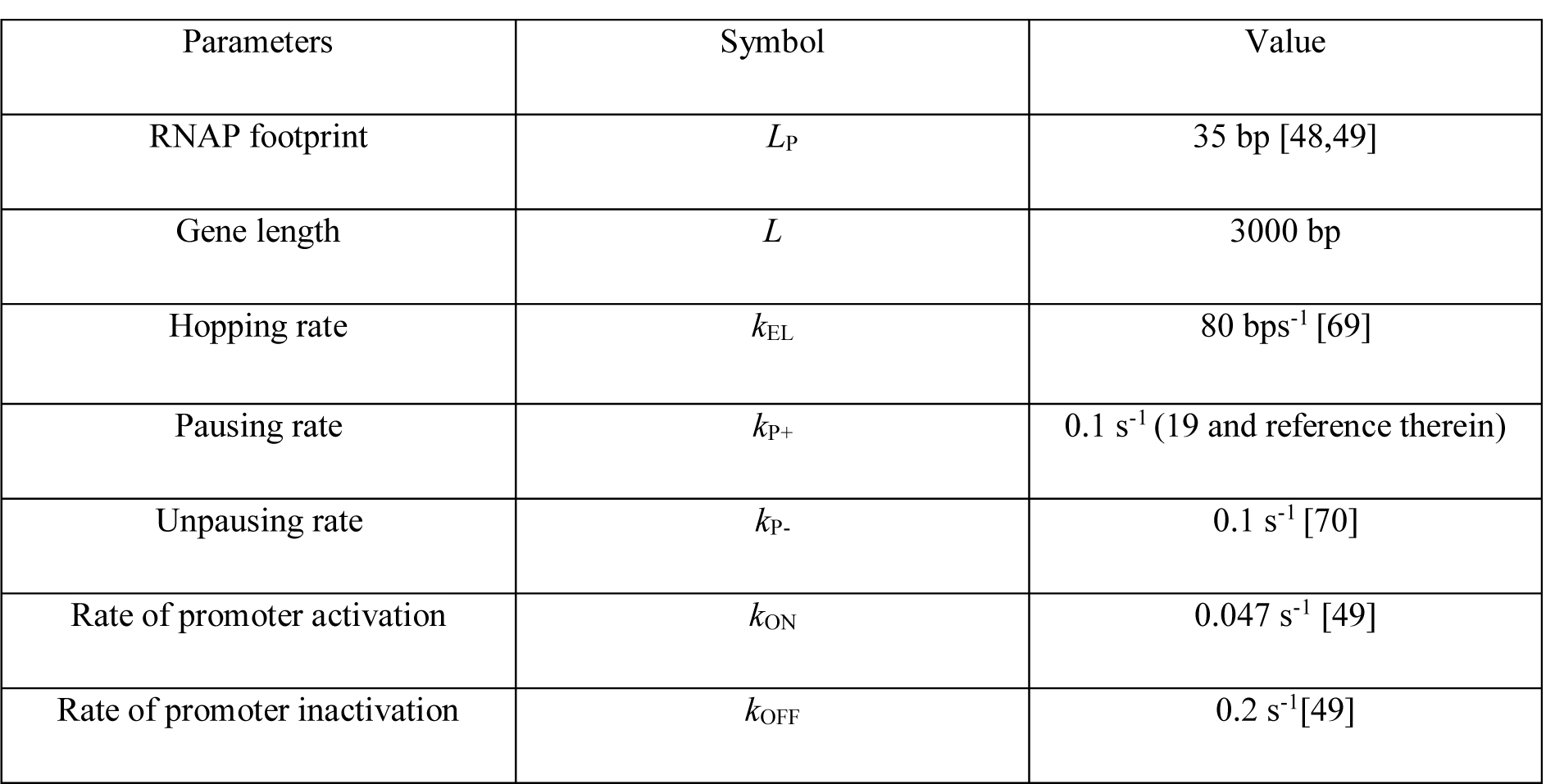
Rate parameters used in Fig. 2.

| Parameters | Symbol | Value |
| --- | --- | --- |
| RNAP footprint | $L_P$ | 35 bp [48,49] |
| Gene length | $L$ | 3000 bp |
| Hopping rate | $k_{EL}$ | 80 bps<sup>-1</sup> [69] |
| Pausing rate | $k_{P+}$ | 0.1 s<sup>-1</sup> (19 and reference therein) |
| Unpausing rate | $k_{P-}$ | 0.1 s<sup>-1</sup> [70] |
| Rate of promoter activation | $k_{ON}$ | 0.047 s<sup>-1</sup> [49] |
| Rate of promoter inactivation | $k_{OFF}$ | 0.2 s<sup>-1</sup> [49] |

**Table 2:** Rate parameters used in Fig. 4.

| Parameters | Symbol | Value | Dataset |
| --- | --- | --- | --- |
| RNAP footprint | $L_P$ | 35 bp [51,69] | All data [51–53] |
| Gene length | $L$ | 6700 bps [51,69] | All data [51–53] |
| Hopping rate | $k_{EL}$ | 60 bps<sup>-1</sup> [51,69] | All data [51–53] |
| Initiation rate<br>(fit to WT data) | $k_{INI}$ | 1 s<sup>-1</sup> | Tongaonkar *et al.* [52] |
|  |  | 1 s<sup>-1</sup> | Schneider *et al.* [53] |
|  |  | 3.5 s<sup>-1</sup> | Claypool *et al.* [51] |
| Rate of promoter<br>activation<br>(fit to WT data) | $k_{ON}$ | 0.01 s<sup>-1</sup> | Tongaonkar *et al.* [52] |
|  |  | 0.015 s<sup>-1</sup> | Schneider *et al.* [53] |
|  |  | 0.04 s<sup>-1</sup> | Claypool *et al.* [51] |
| Rate of promoter<br>inactivation<br>(fit to WT data) | $k_{OFF}$ | 0.012 s<sup>-1</sup> | Tongaonkar *et al.* [52] |
|  |  | 0.013 s<sup>-1</sup> | Schneider *et al.* [53] |
|  |  | 0.012 s<sup>-1</sup> | Claypool *et al.* [51] |

## Results and discussion

### Different models of elongation can be discriminated by their signatures at the level of nascent RNA distribution

The goal of this study is to demonstrate how the cell-to-cell variability of RNAP numbers and inter-polymerase distances can be used to extract dynamical information about the process of transcription elongation *in vivo*. Of particular interest is the dependence of these quantities on the initiation rate; the initiation rate is a readily accessible parameter in experiments, both due to the natural variation in initiation rate, and the ability to systematically alter the rate of initiation by genetic manipulation [47]. The key finding of this paper is that the inclusion of pausing in the model results in qualitatively distinct predictions for the cell-to-cell variability of RNAP numbers and inter-RNAP distances as a function of the initiation rate (Fig. 2) compared to simple elongation (*k*_P+_→ 0). In the following section we discuss the model predictions in the absence of promoter dynamics (a constitutive promoter) by setting the OFF rate (*k*_OFF_) to zero.

**Fig. 2:**
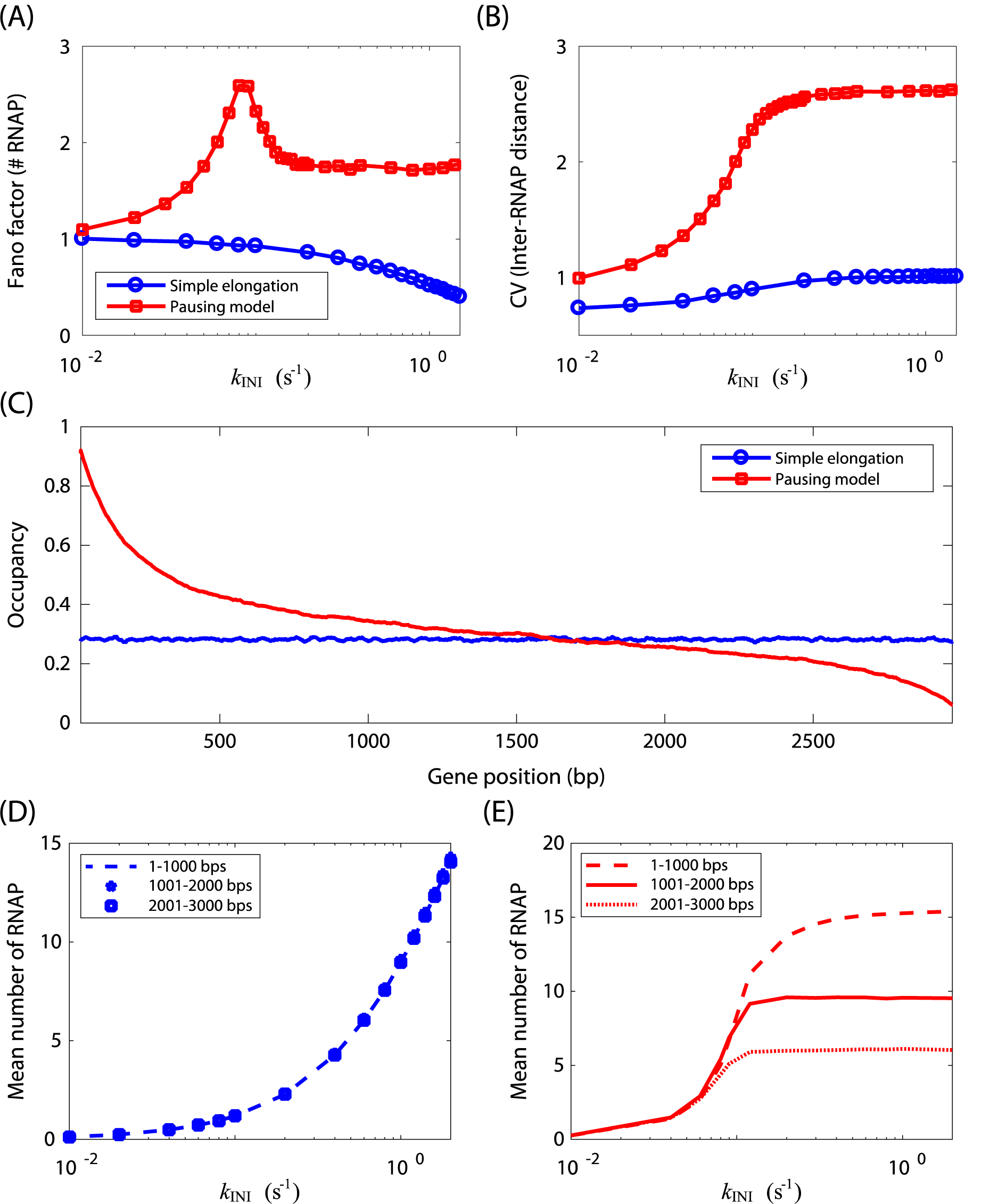
Noise profiles for different models of transcription elongation. (A, B) Using Gillespie simulations [65], we computed the Fano factor of the RNAP number distribution and the coefficient of variation of the inter-polymerase distance distribution along the gene, as a function of the initiation rate of the gene being transcribed, for the two different models of transcription elongation: simple elongation (blue) and pausing model (red). The two different models give qualitatively distinct predictions. To illustrate this point, we use the rates, as shown in Table 1. (C) Probability that a nucleotide on a gene is occupied by an RNAP for simple elongation (*k*_INI_ = 1 s^-1^ shown in blue) and pausing model (*k*_INI_ = 1 s^-1^, *k*_P+_ = 0.1 s^-1^, and *k*_P-_ = 0.1 s^-1^ shown red). (D, E) Mean number of RNAP transcribing a gene for three different regions (1-1000 bp, 1001-2000 bp, and 2001-3000bp) as a function of initiation rate for simple elongation (D) and pausing model (E).

#### Variability in transcribing RNAP numbers

The Fano factor, defined as the ratio of the variance to mean, is a useful measure of the variability in actively transcribing RNAP on a gene because deviation from one indicates departure from a “Poisson” process, *i.e.*, a random process dictated by a single rate-limiting step (initiation, in this case) [48]. As seen in Fig. 2A, when the initiation rate is small, the dynamics of RNAP hopping do not strongly influence the noise profile. The expected distribution of both models is approximately Poissonian and is dictated by the initiation rate. However, for both models, we see a distinct signature in the Fano factor as it deviates from the Poissonian nature of stochastic initiation.

For simple elongation, the Fano factor drops below one as the initiation rate is increased. In this case, collision events between polymerases become more frequent, thus narrowing the polymerase number distribution and the Fano factor decreases below one (blue curve in Fig. 2A). As the density of polymerases on the gene increases, the steric interactions between them order the polymerases along the gene, further reducing variability. In the case of extremely high initiation rate, the polymerases pack the gene with an inter-polymerase distance set by their footprint and thus the variability approaches zero because they are all just hopping along in a line, uniformly spaced. This sub-Poissonian behavior can also be seen by separately observing the mean and variance of the RNAP number distribution as a function of the initiation rate. While both quantities initially increase linearly with the initiation rate, the variance plateaus while the mean continues to rise (see SI Fig. S3). Consequently, Fano factor, being a ratio of the variance and the mean, goes down as the initiation rate is increased.

When pausing is included in the model, the distribution departs from Poisson statistics in a non-monotonic way (red curve in Fig. 2A) due to a combination of two effects: bunch formation driven by pausing and the steric occlusion that reduces variability. When the initiation rate is much smaller than the pausing (*k*_P+_) and unpausing rate (*k*_P-_) of a polymerase molecule, pausing of polymerases on the gene do not impact the RNAP number distribution on a gene [48]. Hence the Fano factor remains close to one. As we increase the initiation rate, the stochastic pausing of a single polymerase molecule on the gene can cause a roadblock. If the lifetime (1/*k*_P-_) of a paused polymerase molecule is long enough (i.e. greater than the time it takes for a polymerase molecule to traverse the average distance between two polymerases) such that polymerase molecules from behind catch up, then bunches of polymerases will form. Bunch formation blocks the procession of polymerases past the stalled polymerase, leading to congestion on the gene which can result in bursts of mRNA production [29,30]. However, some RNAPs do not pause on the gene and bunch formation does not occur, enabling periods without traffic. This leads to a large variability in the time it takes for a polymerase molecule to traverse the gene. The intermediate peak in the Fano factor of RNAP number distribution (see Fig. 2A) corresponds precisely to the peak in variability of travel time (see SI Fig. S12). Snapshots of polymerases on the gene obtained from simulations evidently demonstrate the high variability in bunch formation at intermediate initiation rates (SI Fig. S11). For high initiation rates, bunch formation happens very frequently and typically gives rise to congestion. As a result, at these elevated levels of congestion the variability in the time a polymerase molecule takes to traverse the gene is reduced. Consequently, variability in the polymerase number goes down because polymerases will line up behind roadblocks. Interestingly, the pausing kinetics can also significantly alter the RNAP occupancy along the gene (see Fig. 2C). At high initiation rates, the occupancy is maximum at the start of the gene and gradually decreases towards end of the gene. This feature disappears for low initiation rates and is completely absent for the simple elongation model regardless of initiation rate (Fig. 2C). This is intuitive because downstream jams can contribute to higher density in earlier regions of the gene, however, this is a clear signature of pausing dynamics. As a consequence, separately examining three regions of the gene (1-1000 bp, 1001-2000 bp, and 2001-3000bp), which is experimentally more feasible to examine, reveals that when the initiation rate is small, mean RNAP number in each region is equal (see SI Fig. 2E). But as the initiation rate is increased the number of RNAP in the first one-third of the gene increases and saturates at much higher values compared to the middle one-third, and the end of the gene which has the lowest number of RNAP. Although data of this type is not typically reported, it is experimentally accessible and a strong indicator of the elongation dynamics regardless of the promoter dynamics.

#### Variability in inter-RNAP distances

Next, we seek to unravel the effect of elongation kinetics on the distribution of inter-polymerase distances along a gene. We use coefficient of variation (CV; defined as the ratio of the standard deviation and the mean), a dimensionless quantity, to capture the variability of the inter-polymerase distance distribution along a gene, as shown in Fig. 2B. The CV gives an intuitive interpretation of noise in the inter-polymerase distance distribution; when the initiation rate is much smaller than the hopping rate, the inter-polymerase distances are expected to be exponentially distributed, *i.e.*, CV=1 [49]. The two models of elongation considered above make distinct predictions for the CV, as a function of the initiation rate. It must be noted that the distribution of inter-polymerase distances is restricted by the gene length, *i.e.*, any distances between two polymerases that is greater than the gene length is unphysical and thus the idealized exponential distribution we expect is truncated due to the finite gene length. This feature essentially limits the range of the distribution from zero to *L-L*_P_, where *L* is the gene length and *L*_*P*_ is the RNAP footprint. For the simple elongation model, when the initiation rate is much smaller than the hopping rate i.e., (*k*_EL_/*k*_INI_ ≫ 1) there are few polymerases on the gene at any given instant and inter-polymerase distances on the order of the gene length are frequent. As a result, the gene length restriction impacts the distribution significantly. Consequently, for a finite gene size, the CV is slightly less than one for the simple elongation model (blue line in Fig. 2B). However, as the initiation rate is increased, the gene becomes more densely packed with polymerase and long inter-polymerase distances become rarer. Hence the inter-polymerase distance distribution obtained from simulations captures the first two moments reliably. The CV approaches one as the initiation rate is increased. For the ‘pausing model’, the CV increases to a value greater than one as a function of the initiation rate and later plateaus (see Fig. 2B). Increasing the initiation rate leads to the formation of bunches, as discussed in the previous section. Bunch formation gives rise to two distinct length scales for inter-polymerases distances-one defined by the intra-bunch distances and the other by inter-bunch distances. This bunch formation accounts for the high CV. In fact, this mechanism has been previously suggested as a possible way of generating ‘bursts’ in mRNA production [33]. Importantly, the two models, simple elongation and pausing model, have a distinct imprint on the cell-to-cell variability of inter-polymerase distances that allow us to discriminate between these models from measured distributions of *in vivo* inter-polymerase distance distributions.

The above-mentioned results hold for both models (simple elongation and pausing model) over a wide range of parameter values. For example, choosing a different hopping rate (*k*_EL_ = 40 s^-1^, *k*_EL_ = 20 s^-1^) qualitatively produce the same results as shown in SI Fig. S1, S4. In general, a smaller hopping rate tend to reduce the noise in the RNAP number for both the models. CV of inter-RNAP distance is not affected for the simple elongation model. For the pausing model at higher initiation rate the CV goes down with the hopping rate (see SI Fig. S4). Similarly, varying gene length (*L* = 1000 bp, 5000 bp) also does not affect the results (see SI Fig. S2, S5) qualitatively. However, the frequency or duration of pauses can significantly alter the peak noise as the initiation rate is tuned (see SI Fig. S7, S8).

### Elongation can either enhance or suppress cell-to-cell variability generated by ‘bursty’ transcription initiation

To examine how stochastic initiation and elongation together impact cell-to-cell variability in RNAP numbers and inter-polymerase distances along a gene, we now consider bursty initiation dynamics followed by stochastic elongation. The Fano factor of transcribing RNAP number has been analytically solved when polymerase move deterministically (at a constant speed) along a gene without pausing [23,48,50]. In this case, the RNAPs do not interfere with each other while hopping, and consequently the RNAP number distribution is governed by the promoter dynamics alone. The Fano factor increases linearly as a function of initiation rate for this model, shown as a black dashed line in Fig. 3B (see SI for analytic solution). As such, promoter dynamics alone can result in high cell-to-cell variability in gene expression [14]. However, we find that incorporating stochastic elongation dynamics strongly impacts this relationship.

**Fig. 3:**
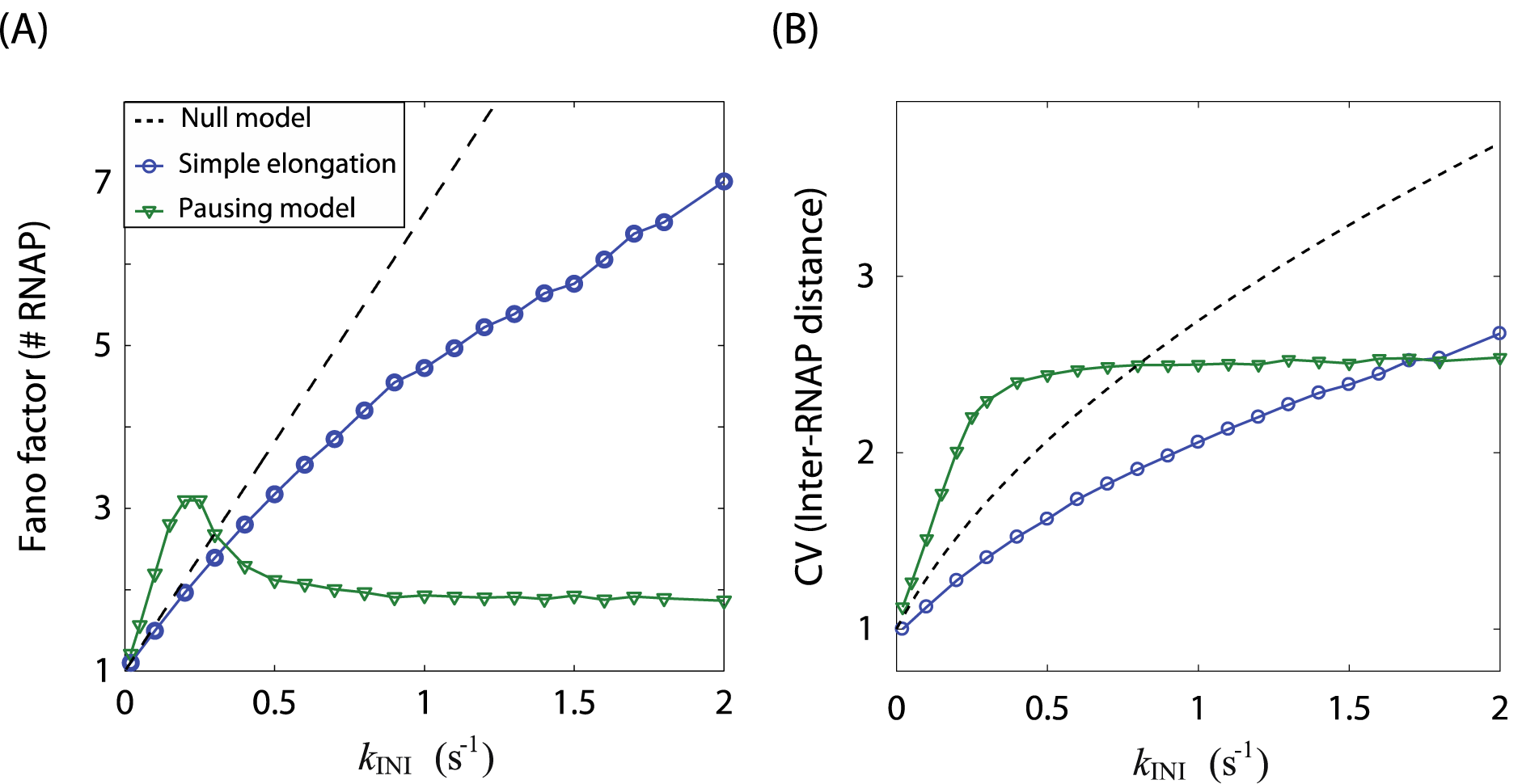
Effect of elongation dynamics on RNAP noise for a bursty promoter. (A) We computed the Fano factor of the nascent RNA number distribution and (B) the coefficient of variation of the inter-polymerase distance distribution as a function of the initiation rate, for the three different models of elongation: on-off promoter dynamics alone (dashed line), simple elongation (blue), pausing model (green) for *k*_P+_ = 0.1 s^-1^ and *k*_P-_ = 0.05 s^-1^. While for the simple elongation Fano factor goes down as the initiation rate is increased, for the pausing model for lower initiation rates an increment in the initiation rate enhances Fano factor. However, for higher initiation rates, any increase in initiation rate decreases Fano factor.

For the bursty initiation model followed by simple elongation, the Fano factor as a function of *k*_INI_ is shown in Fig. 3B (blue line). When the initiation rate is much smaller than the hopping rate, the Fano factor is, once again, dictated by the promoter dynamics. However, as the initiation rate increases, RNAPs start colliding with each other on the gene, resulting in a reduction of the Fano factor compared to promoter dynamics alone; this behavior is similar to the Fano factor reduction observed when initiation occurs at a constant rate (Fig. 2A). As such, we see that simple elongation dynamics can result in dampening some of the variability imparted from the promoter dynamics.

When pausing is included in the elongation dynamics (green line, Fig. 3B), intriguingly, we find that although increasing the rate of initiation initially increases the Fano factor above what is expected from the on-off behavior of the promoter, further ramping up of the initiation rate leads to a suppression of noise generated from the promoter dynamics alone. At low initiation rates, RNAPs that initiate transcription elongation during the promoter’s active period tend to form bunches, which in turn enhances the noise generated from the periodic promoter state switching; for low initiation rates, noise generated at the initiation and elongation levels are additive. Consequently, when the polymerase can pause, the Fano factor for small initiation rates can exceed the Fano factor generated from promoter dynamics alone, as shown in Fig. 3B. However, as the initiation rate is increased further, in the regime where the rate of initiation is comparable or greater than the rate of unpausing of polymerases (*k*_INI_/*k*_P-_ > 1), elongation dynamics govern the RNAP number distribution, in the process disrupting the signature of the initiation dynamics. Hence, the Fano factor of the distribution of RNAP numbers decreases.

We see a similar phenomenon in the CV of the inter-polymerase distances. For the same reasons described above, for small initiation rates the CV of inter-polymerase distances can be higher than the CV expected from the promoter dynamics, as shown in Fig. 3C. As the initiation rate is increased, the inter-polymerase distance distribution is dictated by the elongation dynamics. Consequently, the CV of the inter-polymerase distance distribution gets saturated, as shown in Fig. 3C. These findings hold for a wide range of model parameters, as shown in the SI Fig. S9, S10.

#### Analysis of RNA polymerase number distributions for ribosomal genes in *S. cerevisiae* suggests expression controlled by altering “off rate” of promoter

The theoretical framework developed in the previous sections can be used to extract mechanistic insights about the dynamics of transcription initiation and elongation from *in vivo* measurements of RNAP number distributions. To demonstrate this, we have analyzed three different published data sets of RNAP I (Pol I) distribution on ribosomal RNA (rRNA) genes in budding yeast obtained from electron micrographs (EM) of fixed cells [51–53]. In each dataset, a different perturbation is applied to yeast and the effect on RNAP distributions on the rRNA genes is measured. Ideally, to infer the mechanism of initiation and elongation dynamics one should analyze both the RNAP number and inter-RNAP distance distributions but due to unavailability of the data we look only at the RNAP number distributions in the following sections.

To analyze these data sets, we plot the Fano factor in number of transcribing RNAPs as a function of mean and find that for each data set the studied perturbation leads to a unique response; specifically, one set of data shows a (roughly) constant Fano factor for differing mean levels, while another shows a decrease in Fano factor with the mean, and yet another shows a non-monotonic curve with an intermediate maximum in Fano factor (Fig. 4).

**Fig. 4:**
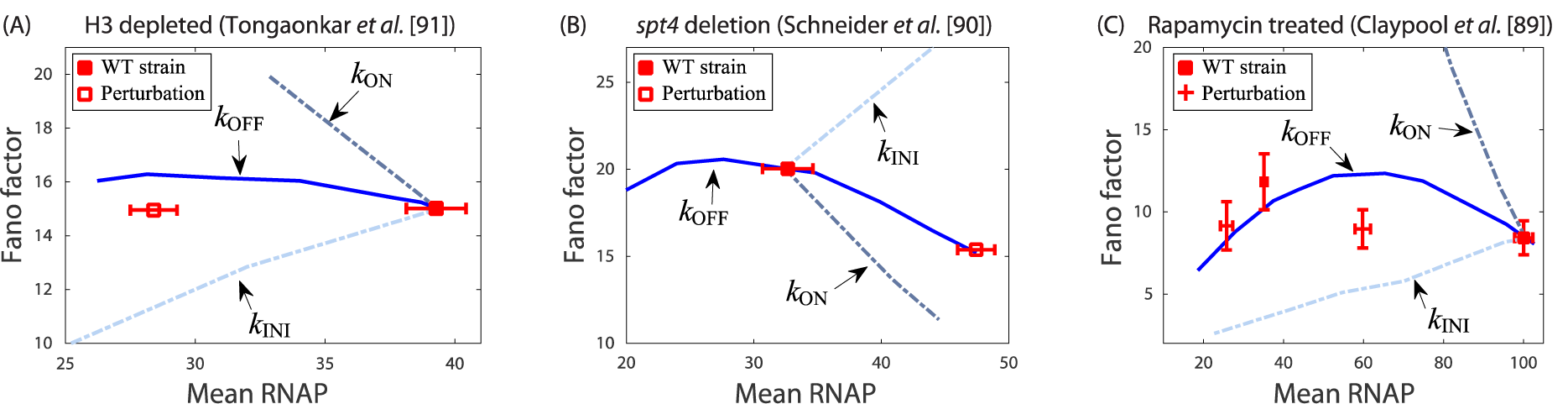
Noise in the RNAP number distributions for ribosomal genes in budding yeast. Fano factor of RNAP number distribution is plotted as a function of mean from the published experimental data [51–53]. For each data set, shown in red squares with error bars, we fit the control stain (WT, filled squares) with the ON-OFF model to extract the parameters (*k*_INI_, *k*_ON,_ and *k*_OFF_; see table 2) and then vary each of them separately keeping the other parameters fixed. Simulation results from varying the initiation rate only (dashed light gray line), promoter activation rate only (dashed dark gray line), and promoter inactivation rate only (solid blue line) are shown for each. Error bars for mean in (A, B) represent standard error of mean (SEM). There are no error bars for Fano factor due to unavailability of raw-data sets. Error bars in (C) represent standard error of mean (SEM), and standard deviation of the Fano factor. The errors in Fano factor are determined by bootstrapping each experimental RNAP number distribution 1000 times and calculating the standard deviation in the Fano factor for those independent bootstrapped data sets.

We find that the data is inconsistent with a constitutive, non-bursty, promoter. This type of promoter with simple elongation predicts a Fano factor of less than or equal to one, which stands in sharp contrast to all these datasets; all measured Fano factors are greater than one. Moreover, although pausing can lead to higher Fano factors and a non-monotonic relationship between mean RNAP numbers and Fano factor, this model is unable to reproduce the large quantitative values exhibited in the data with realistic pausing and unpausing rates (see SI Fig. S14). When pausing is included in the bursty initiation model (as in Fig. 3B), frequent pauses strongly reduce the Fano factor and make it very difficult to match the values for the control strain; as such we find that pausing is not a significant contributor to the RNAP dynamics for these genes. Therefore, below we focus on the model for bursty initiation with simple elongation, although we cannot exclude the possibility that weak pausing exists in the data. As a way to identify the mechanism responsible for the RNAP distribution data we have available, we make predictions for how the Fano factor changes as a function of the mean as any single model parameter is tuned and compare these predictions with the data.

First, for each data set, we extract the parameters characterizing the bursty model (*k*_INI_, *k*_ON_, and *k*_OFF_) using mean RNAP number and the Fano factor of the control strain as constraints, such that the mean RNAP number and the Fano factor is identical to the control wild type strain (defined in the individual descriptions below). Such strategies have been used in simulation by Klump et al [39] to estimate the pause duration in presence/absence of anti-termination complex for ribosomal genes. Other model parameters (RNAP footprint, gene-length, and hopping rate) are directly taken from experimental measurements [11,51] and are kept constant. The fitted values for the control strain of each dataset are listed in Table 2. Upon fitting the WT strains for each dataset, the rates of promoter dynamics we obtain (*k*_ON_ = 0.01, 0.015, 0.04/s, and *k*_OFF_ = 0.012, 0.013, 0.012/s) are in close agreement with the rates reported for Pol II transcription (*k*_ON_ = 0.01/s and *k*_OFF_ = 0.011-0.015/s) [46,54].

Next, we observe the impact of altering any one of these model parameters individually. In each panel of Fig. 4, the solid blue curve corresponds to altering *k*_OFF_, the dark grey curve corresponds to altering *k*_ON_ and the light grey curve corresponds to altering *k*_INI_. Below, we show that for each dataset we can either rule out or highlight which process is being impacted by examining the relationship between the Fano factor and the mean of transcribing RNAP number.

#### Role of histone subunits H3 and H4 in transcription of yeast rRNA genes

In ref. [52], the authors investigate the observation that histone depletion leads to a decrease in transcription of rRNA genes in budding yeast. On one hand, histones are often thought of as barriers that inhibit or slow transcription, however an essential initiating factor, UAF, is often found to contain H3 and H4 implying that the absence of these subunits may cause this decrease in transcription by decreasing the availability of UAF at rRNA genes.

The authors measured RNAP distributions on rRNA genes by examining EM chromatin spreads of two strains: one depleted of H3 and a control strain (WT) with unperturbed H3 levels. They find that a 50% depletion of H3 lowers the average number of polymerases per gene while also keeping the Fano factor roughly constant, these two data points are shown in Fig. 4A. Through various biochemical experiments they indicate that the role of H3 depletion is due to two factors: a decrease in RNAP efficiency (by way of processivity or elongation rate) and periodic “impediments” to initiation. With this in mind, first, we fit the data for the control strain (WT) to find values for the individual rates of the ON-OFF model. Each individual rate of the ON-OFF model is varied over a wide range to get the mean and Fano factor approximately same as the wild type strain (a mean of 39.28 and Fano factor of 15.01, the rates for the control strain is listed in Table 2). Next, we predict how changes in each parameter from the control strain influence the mean and Fano factor (solid and dashed lines in Fig. 4A) and compare the predictions with the data. We find that certain parameters such as modulation of the on rate of the promoter (*k*_ON_) or the initiation rate of the promoter (*k*_INI_) are incompatible with the trend in the data. In fact, the single rate that is compatible with this observation is modulation of the promoter off rate (*k*_OFF_), implying that H3 depletion may cause UAF to be less stable at the promoter, rather than (for instance) decreasing the probability of activation by UAF.

#### Role of Spt4p in transcriptional elongation of rRNA genes

Spt4p and Spt5p are known to form a complex that facilitates transcription elongation in Pol II. Furthermore, it has been shown that this complex competes with an initiation factor, TFE, for the RNAP clamp of Pol II [55].

In ref. [53], the role of Spt4p in transcription by Pol I is investigated by examining chromatin spreads of a control (WT) strain and a strain with *SPT4* deleted. From the examination of chromatin spreads of each of these strains the authors find that the deletion of *SPT4* causes an increase in average polymerase number per gene and a small decrease in Fano factor (data points shown in Fig. 4B). The mechanistic reason for this increase is not explored. First, we find values for the individual rates of the ON-OFF model to match the mean and Fano factor of RNAP number with the control strain (a mean of 32.66 and Fano factor of 20.03, see Table 2) and then varied each rate individually. We find that any alterations in the promoter off rate (*k*_OFF_) or the promoter on rate (*k*_ON_) both influence the noise to mean relationship in a way that is consistent with the data. However, tuning the promoter off rate, shown as solid blue line in Fig. 4B, matches the data points more closely. To distinguish between these two outcomes, we would require more, systematic data points. However, either way we predict that Spt4 is playing a role in the promoter dynamics for Pol I transcription. Naively, we might expect that Spt4 may play a similar role in competing for the clamp of Pol I and thus “turn off” the promoter by out competing an initiation factor.

#### Downregulation of rRNA synthesis in stationary phase

In yeast, gene expression is regulated by different signaling pathways as cells transit from log phase (when cell number grows exponentially in a population) to stationary phase (when cell number on average remains constant in time). In particular, as cells enter the stationary phase, rRNA synthesis decreases [56]. It is also known that chemical treatment of yeast cells (with rapamycin) inhibits the functions of Tor signaling pathway, and also inhibits transcription of rRNA genes, leading to a state that mimics the stationary phase.

Using the above systematic perturbation of a control yeast strain in the log phase, Claypool *et al.* [51] investigated how the Tor signaling pathway affects rRNA transcription. The authors showed that a pre-initiation complex, consisting of Pol I and a specific transcription factor (Rrn3p), is almost absent in the chemically perturbed cells, and also in the stationary phase. This reduction in the pre-initiation complex thus explains the down-regulation of rRNA synthesis and suggests that Tor signaling pathway acts on the step of transcription initiation. To quantify the down-regulation of rRNA synthesis, the authors calculated the mean and variance of RNAP number distributions from EM images of the fixed cells. This was done for four different cases: (i) cells in log phase (used as a control), (ii) cells treated with rapamycin for 10 min, (iii) cells treated with rapamycin for 30 min, and (iii) cells in stationary phase. When the chemical inhibition of Tor pathway is systematically intensified, the mean RNAP number per gene decreases from the control and reaches a minimum in the stationary phase, as shown in Fig. 4B. Surprisingly, the EM images showed that when the cells transit from the log phase to the stationary phase, a very different pattern of polymerase occupancy of the rRNA genes emerges. While the mean number of RNAP molecules along each gene decreases, the Fano factor of the RNAP distribution shows a non-monotonic behavior showing an intermediate peak.

Upon fitting our model to the control strain (a mean of 110.06 and Fano factor of 8.41, see Table 2) we further check that the extracted parameters correctly reproduce the measured range of transcription rate for the control strain [51] (the experimental value is 0.8/s – 1.1/s per gene, while the calculated value from the simulations is 0.9/s). While modulating any of the three parameters can produce a downregulation of the mean number of RNAPs on the ribosomal genes, only an alteration in the rate of promoter-inactivation (*k*_OFF_) matches the trend of the data (see Fig. 4C). Changing the rate of initiation (*k*_INI_) and promoter activation (*k*_ON_) either lead to a decrease or an increase in the Fano factor as functions of the mean (dashed lines in Fig. 4C). A change in the rate of promoter inactivation alone predicts a non-monotonic behavior of the Fano factor as a function of the mean: The Fano factor first increases to a maximum and then decreases, shown as solid blue line in Fig. 4C. Assuming only one of these rates is tuned, our results thus signify that yeast cells most likely regulate the inactivation rate of the bursty promoters as they approach the stationary phase from the log phase. That being said, more systematic data, particularly between 40 and 80 polymerases per gene, would help to further substantiate this finding. Regardless, our analysis demonstrates how useful mechanistic information about the initiation and elongation dynamics can be extracted from the experimental data by employing a mathematical framework akin to that demonstrated here.

## Discussion

Regulation of gene expression at the level of transcription has been a topic of intense inquiry over the last few decades [57,58]. Consequently, the dynamics of initiation has received a lot of attention from researchers and led to a deepening of understanding of how it’s regulated. This view though is getting challenged in the light of recent experimental findings; the events downstream of transcription initiation, such as RNAP pausing along the gene play significant roles in regulating RNA production. Hence it is imperative to catalog how the different steps in transcription contribute to the regulation of gene expression in order to develop a comprehensive understanding. In this paper, we examine a common model of transcription initiation and elongation dynamics in order to predict the impact of these processes on nascent RNA (RNAs in the process of being synthesized by the RNAP molecule) distributions. Recent experimental advancements bestow a unique opportunity to make progress in this direction. Single molecule FISH measurements and EM images of transcribing RNAPs provide the distribution of RNAP molecules along a gene across an isogenic population. Motivated by these advances, we utilize a kinetic model of transcription to connect mechanisms of transcription to this type of experimental data. The models of transcription initiation and elongation considered here provide a route to delineate the impact of elongation on the distributions of RNAP numbers and inter-polymerase distances. In particular, we consider two different models of transcription elongation that incorporate two broad classes of elongation mechanisms and show that they make qualitatively distinct predictions for the distributions of RNAP numbers and inter-polymerase distances, which in turn allows us to discern them. For bursty initiation followed by simple elongation cell-to-cell variability at the RNAP numbers, generated at the level of the initiation get suppressed due to the collision between RNAP molecules on the gene. Intriguingly though for the ‘pausing model’ elongation dynamics can have a non-monotonic influence on RNAP number distribution, as we increase the initiation rate. While in the beginning the noise is determined primarily by the initiation dynamics, a further increase in initiation rate leads to a regime where the cell-to-cell variability due to initiation and elongation dynamics are additive. On the other hand, for an even higher initiation rate, RNAP number distribution is primarily dominated by the elongation dynamics.

Armed with these simulation results, we analyzed three available RNAP number distribution data for ribosomal genes in budding yeast (*S. cerevisiae*) [51–53]. Our analysis reveals that when this data is interpreted through a simple ON-OFF model of transcription, the mechanistic effect of each experiment can be identified, or at least narrowed down. For instance, in all cases the Fano factor is much greater than one, immediately ruling out such simple models as stochastic initiation (a promoter that does not have an off-state) followed by very efficient elongation by polymerase *i.e.* simple elongation or insignificant pausing events. In all cases, we find that a bursty initiation model followed by simple elongation can explain the data well. Our data analysis finds that for each case perturbations caused the yeast cells to tune the rate of promoter-inactivation to modify rRNA synthesis rates. However, importantly, our model considers only intrinsic sources of noise and does not explicitly consider extrinsic sources that would result from variability in the cellular environment of individual cells [59–61]. Such sources of noise can be included in simulations by treating the rates of, for instance, initiation or promoter switching as stemming from a distribution of possible values that reflect differences in cellular concentrations of the responsible proteins.

While using noise in gene expression has been a successful strategy in disentangling the molecular mechanism of transcription initiation [7,9,62–64], our theoretical attempt can provide a further leap in developing a comprehensive understanding of how initiation and elongation together affect nascent RNA distribution.

## Methods

### Computational method

We implement the simulations using Gillespie’s algorithm [65] for stochastic reaction systems in C/Matlab. The gene template is assumed to be a one-dimensional lattice of length L, where each lattice site represents one base-pair of the gene. Transcription initiation starts by loading RNAP on the gene template with rate *k*_INI_, if the first 35 bp of the gene, treated as promoter region, is not already occupied by another RNAP. The RNAP then stochastically move forward, a one base-pair step, with rate *k*_EL_. The forward hopping occurs only if the lattice site is not occluded by another RNAP. It is assumed that at each lattice site a transcribing RNAP pauses with a probability determined by the pausing rate *k*_P+_. A paused RNAP switches back to active state stochastically with a rate *k*_P-_ and continue to hop forward with rate *k*_EL_. For simple elongation model the probability of pausing is set to zero. Termination of RNAP occurs when a forward step is taken at the last base of the gene. For the bursty initiation model, the promoter switches back and forth between an ON state and an OFF state with rate *k*_OFF_ and *k*_ON_, respectively. Transcription initiation occurs only when the promoter is in ON state. Each simulation is run for sufficiently long time (∽10^5^ s) to reach the steady state. Typically, for the rates used in the paper the steady state is achieved in 10^2^-10^4^ s. Data for steady state distributions (number of RNAP molecules on the gene template and the inter-RNAP distances) are then recorded by sampling over time with a time interval (*T*_S_) long enough for the slowest reaction to occur 20 times on an average (*T*_S_ = 20 / rate for slowest reaction). Fano factor and CV is calculated using at least 30,000 data points for each set of parameters. The rates of the reactions are kept constant for any given simulation, i.e., the cell-to-cell variability comes only from the stochasticity of the reaction steps and not from the extrinsic sources such as population of cells having different initiation rate, hopping rate etc.

### Parameter selection

Model parameters used in the simulation are listed in table 1 and 2 and are taken from previously reported papers. The pausing frequency (*k*_P+_) and pause duration (1/*k*_P-_) has been measured *in vitro* using single-molecule experiments [43,66]. Experiments have revealed that RNAP exhibit both short pauses (majority of the pauses lasting less than 10 s with a frequency of 0.07-0.15/s) and longer pauses (duration of ∽ 20 s). The longer pauses are found to be strongly sequence dependent and associated with the backtracking of RNAPs. The ubiquitous short pauses on the other hand show a weak sequence dependence. In our simulation we have checked that if RNAP pauses at 1 site per 100 bps gives similar results when pausing occur at every nucleotide (the frequency of pausing being tuned to get similar number of pauses during elongation). Hence to keep the model simple we consider that the pauses are sequence independent. In our simulation we have used a pausing rate in a range 0.1-1/s and unpausing rate in a rage 0.1-1/s (i.e. pause duration of 1-10 s). The initiation rate, *k*_INI_, is varied from 0.01-2/s. We have used elongation rate, *k*_EL_= 80/s throughout the paper unless specified. We chose the rate such that the average velocity of elongating RNAPs are within the range of 10-80 bps/s [67,68]. We have used *k*_ON_ = 0.047/s and *k*_OFF_ = 0.2/s to incorporate promoter dynamics in Fig. 3. These rates were previously estimated by Choubey *et. al.* [49] from EM images of transcribing RNAP molecules attached to ribosomal genes. For the data analysis of the ribosomal genes the on and off rates are estimated by fitting the mean and the Fano of transcribing RNAPs and are listed in Table 2.

### Analysis of the experimental data

To test our model predictions, we analyze experimental data for the RNAP number distribution from three different studies [51–53]. For the dataset from Claypool *et al.* [51], the number of polymerase counted for individual genes was provided by the authors. From this raw data, we corrected these measurements by adding zero counts to each data set corresponding to the fraction of inactive genes. Error bars in both mean and Fano factor represent the standard deviation of 1000 independent, resampled data sets with replacement obtained using the method of Bootstrapping in Matlab.

Due to unavailability of raw data for other two data sets analyzed [90,91], the Fano factor is recalculated to correct for the inactive gene fraction by incorporating the following simple algebraic calculation. We assume that for a given dataset without zeros, *x*_*i*_ (*i=*1:*n*), the mean and the variance (square of the standard deviation) are *u*_*0*_ and *var*_*0*_, respectively. These values are reported in the References [52,53]. Now, to correct for the number of inactive genes, denoted by letter *m*, we use the following equations

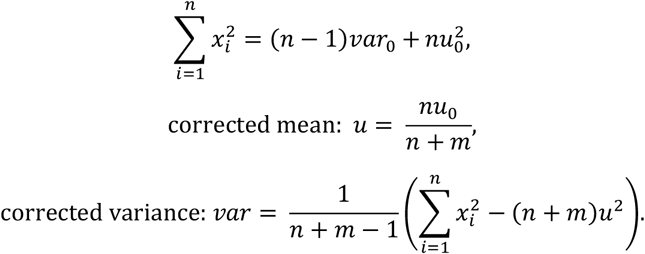

Error bars in the mean of these measurements is the standard error, however because we do not have the raw data, we cannot compute an error bar for the Fano factor of these data. The final corrected mean and Fano factor are listed in table 3 for each study.

**Table 3:** Correcting for inactive gene copies in experimental datasets.

| Strain | $u_0$ | $var_0$ | Active/Inactive<br>(n/m) | Corrected<br>Mean ($u$) | Corrected<br>Fano ($var / u$) |
| --- | --- | --- | --- | --- | --- |
| WT (177 copy) [53] | 46 | 302.76 | 29/71 | 32.66 | 20.03 |
| *spt4Δ* [53] | 59.3 | 201.64 | 20/80 | 47.44 | 15.37 |
| WT [52] | 49 | 256 | 23/93 | 39.28 | 15.01 |
| H3 depleted [52] | 35 | 289 | 10/43 | 28.4 | 14.95 |
| WT (40 copy) [51] | - | - | - | 100.06 | 8.41 |
| 10 min rapamycin (40 copy) [51] | - | - | 2/233 | 59.81 | 8.96 |
| 30 min rapamycin (40 copy) [51] | - | - | 2/202 | 35.15 | 11.82 |
| Post-log phase (40 copy) [51] | - | - | 22/229 | 25.75 | 9.15 |

## Supplementary Data

Supplementary Data are available online.

## Acknowledgements

We wish to thank Sarah French for kindly providing the raw data set from ref. [51]. We also wish to thank Jane Kondev, Alvaro Sanchez for years of stimulating discussions and shared thoughts about how transcriptional dynamics impacts noise. Md. ZA would like to thank Sunil Guharajan for helpful discussions. DD would like to thank Supravat Dey for the initial discussions on the data analysis. Research reported in this publication was supported by the National Institutes of Health under award number R35 GM128797.

## References

1. Shandilya J, Roberts SGE. The transcription cycle in eukaryotes: From productive initiation to RNA polymerase II recycling. Biochim Biophys Acta BBA - Gene Regul Mech. 2012;1819: 391–400. doi:10.1016/j.bbagrm.2012.01.010

2. Svejstrup JQ. The RNA polymerase II transcription cycle: cycling through chromatin. Biochim Biophys Acta BBA - Gene Struct Expr. 2004;1677: 64–73. doi:10.1016/j.bbaexp.2003.10.012

3. Mayer A, Landry HM, Churchman LS. Pause & go: from the discovery of RNA polymerase pausing to its functional implications. Curr Opin Cell Biol. 2017;46: 72–80. doi:10.1016/j.ceb.2017.03.002

4. Gandhi SJ, Zenklusen D, Lionnet T, Singer RH. Transcription of functionally related constitutive genes is not coordinated. Nat Struct Mol Biol. 2011;18: 27–34. doi:10.1038/nsmb.1934

5. Churchman LS, Weissman JS. Nascent transcript sequencing visualizes transcription at nucleotide resolution. Nature. 2011;469. doi:10.1038/nature09652

6. Larson DR, Zenklusen D, Wu B, Chao JA, Singer RH. Real-Time Observation of Transcription Initiation and Elongation on an Endogenous Yeast Gene. Science. 2011;332: 475–478. doi:10.1126/science.1202142

7. Sanchez A, Garcia HG, Jones D, Phillips R, Kondev J. Effect of Promoter Architecture on the Cell-to-Cell Variability in Gene Expression. PLoS Comput Biol. 2011;7: e1001100. doi:10.1371/journal.pcbi.1001100

8. Kepler TB, Elston TC. Stochasticity in transcriptional regulation: origins, consequences, and mathematical representations. Biophys J. 2001;81: 3116–3136. doi:10.1016/S0006-3495(01)75949-8

9. Munsky B, Neuert G, Oudenaarden A van. Using Gene Expression Noise to Understand Gene Regulation. Science. 2012;336: 183–187. doi:10.1126/science.1216379

10. Gotta SL, Miller OL, French SL. rRNA transcription rate in Escherichia coli. J Bacteriol. 1991;173: 6647–6649.

11. Voulgaris J, French S, Gourse RL, Squires C, Squires CL. Increased rrn gene dosage causes intermittent transcription of rRNA in Escherichia coli. J Bacteriol. 1999;181: 4170–4175.

12. Condon C, French S, Squires C, Squires CL. Depletion of functional ribosomal RNA operons in Escherichia coli causes increased expression of the remaining intact copies. EMBO J. 1993;12: 4305–4315.

13. El Hage A, French SL, Beyer AL, Tollervey D. Loss of Topoisomerase I leads to R-loop-mediated transcriptional blocks during ribosomal RNA synthesis. Genes Dev. 2010;24: 1546–1558. doi:10.1101/gad.573310

14. Zenklusen D, Larson DR, Singer RH. Single-RNA counting reveals alternative modes of gene expression in yeast. Nat Struct Mol Biol. 2008;15: 1263–1271. doi:10.1038/nsmb.1514

15. Larson DR, Singer RH, Zenklusen D. A single molecule view of gene expression. Trends Cell Biol. 2009;19: 630–637. doi:10.1016/j.tcb.2009.08.008

16. Castelnuovo M, Rahman S, Guffanti E, Infantino V, Stutz F, Zenklusen D. Bimodal expression of PHO84 is modulated by early termination of antisense transcription. Nat Struct Mol Biol. 2013;20: 851–858. doi:10.1038/nsmb.2598

17. Anderson SJ, Sikes ML, Zhang Y, French SL, Salgia S, Beyer AL, et al. The transcription elongation factor Spt5 influences transcription by RNA polymerase I positively and negatively. J Biol Chem. 2011; jbc.M110.202101. doi:10.1074/jbc.M110.202101

18. Quan S, Zhang N, French S, Squires CL. Transcriptional Polarity in rRNA Operons of Escherichia coli nusA and nusB Mutant Strains. J Bacteriol. 2005;187: 1632–1638. doi:10.1128/JB.187.5.1632-1638.2005

19. Kepler TB, Elston TC. Stochasticity in transcriptional regulation: origins, consequences, and mathematical representations. Biophys J. 2001;81: 3116–3136. doi:10.1016/S0006-3495(01)75949-8

20. Kaern M, Elston TC, Blake WJ, Collins JJ. Stochasticity in gene expression: from theories to phenotypes. Nat Rev Genet. 2005;6: 451–464. doi:10.1038/nrg1615

21. Raj A, Oudenaarden A van. Single-Molecule Approaches to Stochastic Gene Expression. Annu Rev Biophys. 2009;38: 255–270. doi:10.1146/annurev.biophys.37.032807.125928

22. Jones DL, Brewster RC, Phillips R. Promoter architecture dictates cell-to-cell variability in gene expression. Science. 2014;346: 1533–1536. doi:10.1126/science.1255301

23. Choubey S. Nascent RNA kinetics: Transient and steady state behavior of models of transcription. Phys Rev E. 2018;97: 022402. doi:10.1103/PhysRevE.97.022402

24. Sanchez A, Choubey S, Kondev J. Regulation of Noise in Gene Expression. Annu Rev Biophys. 2013;42: 469–491. doi:10.1146/annurev-biophys-083012-130401

25. Das D, Dey S, Brewster RC, Choubey S. Effect of transcription factor resource sharing on gene expression noise. PLoS Comput Biol. 2017;13: e1005491. doi:10.1371/journal.pcbi.1005491

26. Li G-W, Xie XS. Central dogma at the single-molecule level in living cells. Nature. 2011;475: 308–315. doi:10.1038/nature10315

27. Cai L, Friedman N, Xie XS. Stochastic protein expression in individual cells at the single molecule level. Nature. 2006;440: 358–362. doi:10.1038/nature04599

28. Jia T, Kulkarni RV. Intrinsic noise in stochastic models of gene expression with molecular memory and bursting. Phys Rev Lett. 2011;106: 058102.

29. Rajala T, Häkkinen A, Healy S, Yli-Harja O, Ribeiro AS. Effects of Transcriptional Pausing on Gene Expression Dynamics. PLOS Comput Biol. 2010;6: e1000704. doi:10.1371/journal.pcbi.1000704

30. Kim S, Jacobs-Wagner C. Effects of mRNA Degradation and Site-Specific Transcriptional Pausing on Protein Expression Noise. Biophys J. 2018;114: 1718–1729. doi:10.1016/j.bpj.2018.02.010

31. Platini T, Jia T, Kulkarni RV. Regulation by small RNAs via coupled degradation: Mean-field and variational approaches. Phys Rev E. 2011;84: 021928. doi:10.1103/PhysRevE.84.021928

32. Jia T, Kulkarni RV. Post-transcriptional regulation of noise in protein distributions during gene expression. Phys Rev Lett. 2010;105: 018101.

33. Singh A, Bokes P. Consequences of mRNA Transport on Stochastic Variability in Protein Levels. Biophys J. 2012;103: 1087–1096. doi:10.1016/j.bpj.2012.07.015

34. Huh D, Paulsson J. Random partitioning of molecules at cell division. Proc Natl Acad Sci U S A. 2011;108: 15004–15009. doi:10.1073/pnas.1013171108

35. Huh D, Paulsson J. Non-genetic heterogeneity from random partitioning at cell division. Nat Genet. 2011;43: 95–100. doi:10.1038/ng.729

36. Schmidt U, Basyuk E, Robert M-C, Yoshida M, Villemin J-P, Auboeuf D, et al. Real-time imaging of cotranscriptional splicing reveals a kinetic model that reduces noise: implications for alternative splicing regulation. J Cell Biol. 2011;193: 819–829. doi:10.1083/jcb.201009012

37. Melamud E, Moult J. Stochastic noise in splicing machinery. Nucleic Acids Res. 2009;37: 4873–4886. doi:10.1093/nar/gkp471

38. Dobrzyński M, Bruggeman FJ. Elongation dynamics shape bursty transcription and translation. Proc Natl Acad Sci. 2009;106: 2583–2588. doi:10.1073/pnas.0803507106

39. Klumpp S, Hwa T. Stochasticity and traffic jams in the transcription of ribosomal RNA: Intriguing role of termination and antitermination. Proc Natl Acad Sci. 2008;105: 18159–18164. doi:10.1073/pnas.0806084105

40. Klumpp S. Pausing and Backtracking in Transcription Under Dense Traffic Conditions. J Stat Phys. 2011;142: 1252–1267. doi:10.1007/s10955-011-0120-3

41. Voliotis M, Cohen N, Molina-París C, Liverpool TB. Fluctuations, pauses, and backtracking in DNA transcription. Biophys J. 2008;94: 334–348. doi:10.1529/biophysj.107.105767

42. Tinoco I, Bustamante C. The effect of force on thermodynamics and kinetics of single molecule reactions. Biophys Chem. 2002;101–102: 513–533.

43. Forde NR, Izhaky D, Woodcock GR, Wuite GJL, Bustamante C. Using mechanical force to probe the mechanism of pausing and arrest during continuous elongation by Escherichia coli RNA polymerase. Proc Natl Acad Sci U S A. 2002;99: 11682–11687. doi:10.1073/pnas.142417799

44. van den Berg AA, Depken M. Crowding-induced transcriptional bursts dictate polymerase and nucleosome density profiles along genes. Nucleic Acids Res. 2017;45: 7623–7632. doi:10.1093/nar/gkx513

45. Mitarai N, Dodd IB, Crooks MT, Sneppen K. The generation of promoter-mediated transcriptional noise in bacteria. PLoS Comput Biol. 2008;4: e1000109. doi:10.1371/journal.pcbi.1000109

46. Tantale K, Mueller F, Kozulic-Pirher A, Lesne A, Victor J-M, Robert M-C, et al. A single-molecule view of transcription reveals convoys of RNA polymerases and multi-scale bursting. Nat Commun. 2016;7: ncomms12248. doi:10.1038/ncomms12248

47. Brewster RC, Jones DL, Phillips R. Tuning promoter strength through RNA polymerase binding site design in Escherichia coli. PLoS Comput Biol. 2012;8: e1002811. doi:10.1371/journal.pcbi.1002811

48. Choubey S, Kondev J, Sanchez A. Deciphering Transcriptional Dynamics In Vivo by Counting Nascent RNA Molecules. PLOS Comput Biol. 2015;11: e1004345. doi:10.1371/journal.pcbi.1004345

49. Choubey S, Kondev J, Sanchez A. Distribution of Initiation Times Reveals Mechanisms of Transcriptional Regulation in Single Cells. Biophys J. 2018;114: 2072–2082. doi:10.1016/j.bpj.2018.03.031

50. Zoller B, Little SC, Gregor T. Diverse Spatial Expression Patterns Emerge from Unified Kinetics of Transcriptional Bursting. Cell. 2018;175: 835–847.e25. doi:10.1016/j.cell.2018.09.056

51. Claypool JA, French SL, Johzuka K, Eliason K, Vu L, Dodd JA, et al. Tor Pathway Regulates Rrn3p-dependent Recruitment of Yeast RNA Polymerase I to the Promoter but Does Not Participate in Alteration of the Number of Active Genes. Mol Biol Cell. 2004;15: 946–956. doi:10.1091/mbc.E03-08-0594

52. Tongaonkar P, French SL, Oakes ML, Vu L, Schneider DA, Beyer AL, et al. Histones are required for transcription of yeast rRNA genes by RNA polymerase I. Proc Natl Acad Sci. 2005;102: 10129–10134. doi:10.1073/pnas.0504563102

53. Schneider DA, French SL, Osheim YN, Bailey AO, Vu L, Dodd J, et al. RNA polymerase II elongation factors Spt4p and Spt5p play roles in transcription elongation by RNA polymerase I and rRNA processing. Proc Natl Acad Sci. 2006;103: 12707–12712. doi:10.1073/pnas.0605686103

54. Fritzsch C, Baumgärtner S, Kuban M, Steinshorn D, Reid G, Legewie S. Estrogen-dependent control and cell-to-cell variability of transcriptional bursting. Mol Syst Biol. 2018;14: e7678. doi:10.15252/msb.20177678

55. Grohmann D, Nagy J, Chakraborty A, Klose D, Fielden D, Ebright RH, et al. The initiation factor TFE and the elongation factor Spt4/5 compete for the RNAP clamp during transcription initiation and elongation. Mol Cell. 2011;43: 263–274. doi:10.1016/j.molcel.2011.05.030

56. Sandmeier JJ, French S, Osheim Y, Cheung WL, Gallo CM, Beyer AL, et al. RPD3 is required for the inactivation of yeast ribosomal DNA genes in stationary phase. EMBO J. 2002;21: 4959–4968. doi:10.1093/emboj/cdf498

57. Cooper GM. Regulation of Transcription in Eukaryotes. 2000; Available:https://www.ncbi.nlm.nih.gov/books/NBK9904/

58. Ptashne M, Gann A. Transcriptional activation by recruitment. Nature. 1997;386: 569–577. doi:10.1038/386569a0

59. Elowitz MB, Levine AJ, Siggia ED, Swain PS. Stochastic gene expression in a single cell. Science. 2002;297: 1183–1186. doi:10.1126/science.1070919

60. Swain PS, Elowitz MB, Siggia ED. Intrinsic and extrinsic contributions to stochasticity in gene expression. Proc Natl Acad Sci U S A. 2002;99: 12795–12800. doi:10.1073/pnas.162041399

61. Hilfinger A, Paulsson J. Separating intrinsic from extrinsic fluctuations in dynamic biological systems. Proc Natl Acad Sci U S A. 2011;108: 12167–12172. doi:10.1073/pnas.1018832108

62. Kar G, Kim JK, Kolodziejczyk AA, Natarajan KN, Triglia ET, Mifsud B, et al. Flipping between Polycomb repressed and active transcriptional states introduces noise in gene expression. Nat Commun. 2017;8: 36. doi:10.1038/s41467-017-00052-2

63. Raser JM, O’Shea EK. Noise in Gene Expression: Origins, Consequences, and Control. Science. 2005;309: 2010–2013. doi:10.1126/science.1105891

64. Jones DL, Brewster RC, Phillips R. Promoter architecture dictates cell-to-cell variability in gene expression. Science. 2014;346: 1533–1536. doi:10.1126/science.1255301

65. Gillespie DT. Exact stochastic simulation of coupled chemical reactions. J Phys Chem. 1977;81: 2340–2361. doi:10.1021/j100540a008

66. Adelman K, La Porta A, Santangelo TJ, Lis JT, Roberts JW, Wang MD. Single molecule analysis of RNA polymerase elongation reveals uniform kinetic behavior. Proc Natl Acad Sci U S A. 2002;99: 13538–13543. doi:10.1073/pnas.212358999

67. Epshtein V, Nudler E. Cooperation between RNA polymerase molecules in transcription elongation. Science. 2003;300: 801–805. doi:10.1126/science.1083219

68. Vogel U, Jensen KF. The RNA chain elongation rate in Escherichia coli depends on the growth rate. J Bacteriol. 1994;176: 2807–2813.

69. French SL, Osheim YN, Cioci F, Nomura M, Beyer AL. In exponentially growing Saccharomyces cerevisiae cells, rRNA synthesis is determined by the summed RNA polymerase I loading rate rather than by the number of active genes. Mol Cell Biol. 2003;23: 1558–1568.

70. Klumpp S, Hwa T. Stochasticity and traffic jams in the transcription of ribosomal RNA: Intriguing role of termination and antitermination. Proc Natl Acad Sci. 2008;105: 18159–18164. doi:10.1073/pnas.0806084105

