## Supplementary Information for "Probing mechanisms of transcription elongation through cell-to-cell variability of RNA polymerase"


**Analytical solution for the promoter dynamics of transcriptional regulation**

One of the key contributions of this paper involves the exploration of how transcription initiation and elongation together impact the noise in RNAP numbers and inter-polymerase distances. We consider the bursty initiation model followed by 1) simple elongation and 2) pausing. We compare the noise generated from these models to the promoter dynamics alone, where bursty initiation is followed by deterministic elongation. This allows us to decipher how stochastic elongation kinetics impacts the noise in RNAP numbers and inter-polymerase distances generated through initiation dynamics. Previous studies have analytically obtained the Fano factor of the RNAP numbers [1,2] and CV of the inter-polymerase distances [3]. The Fano factor of the polymerase numbers is given by,

$$Fano\ factor = \ 1 + \frac{2k_{INI}k_{OFF}}{(k_{ON}+k_{OFF})^2} + \frac{2k_{INI}k_{OFF}}{T(k_{ON}+k_{OFF})^3}[\exp\{-(k_{ON}+k_{OFF})T\} - 1].$$

Here $k_{\mathrm{OFF}}$ is the rate of promoter switching from the active to the inactive state, $k_{\mathrm{ON}}$ is the rate of promoter switching from the inactive to the active state, $k_{\mathrm{INI}}$ is the rate of transcription initiation from the active state, and the time of elongation is $T=L/k_{\mathrm{EL}}$.

The CV of the inter-polymerase distances is given by,

$$CV = \frac{[(k_{ON}+k_{OFF})^2 + 2k_{OFF}k_{INI}]^{\frac{1}{2}}}{(k_{ON}+k_{OFF})}.$$

We use these formulas to explore how the Fano factor and the CV change as functions of initiation rate i.e. $k_{\mathrm{INI}}$.

**Sequence dependent pausing**

In this section, we show that sequence dependent pausing does not alter our conclusion in the main text where the assumption is that pauses occur and resolve with equal probability at every base. To incorporate sequence specific pausing, we now assume that RNAP molecules can pause at every 100 bps along the gene such that there are $L/100$ pausing sites on the gene (choosing a random pausing site every 100 bps results in the same conclusions). Again, the rate of switching from active to paused state and back to active state are $k_{\mathrm{P+}}$ and $k_{\mathrm{P-}}$ respectively and is same for each of these sites. To match the average spatial frequency of pauses we use a pausing rate 100 times higher than the pausing at random sites. We find that the qualitative features with and without site specific pausing are similar and are in sharp contrast to the simple elongation model (see Fig. S13). The quantitative values, however, show variation and depend on specific details of the pausing sites.

### These authors contributed equally to the paper as first authors.

**Reactions describing the model**

Let $L$ be the length of the gene and $n$ the index of a given base (nucleotide) on the gene. The occupied and free bases are denoted by $O_n$ and $U_n$ respectively, where the subscript n goes from 1 to L. The first $L_p$ bps is treated as the promoter region where $L_p$ also denotes the of the size of RNAP molecules such that if an RNAP molecules is on a gene its position is given by $O_{n:n+L_p-1}$. The promoter switches between an ON state and OFF state with rates $k_{OFF}$ and $k_{ON}$, respectively and is given by

$$U_{1:L_p}^{ON} \xrightarrow{k_{OFF}} U_{1:L_p}^{OFF} \;,$$
$$U_{1:L_p}^{OFF} \xrightarrow{k_{ON}} U_{1:L_p}^{ON} \;.$$

When the promoter is active and unoccupied by previously initiated RNAP, initiation occur with rate $k_{INI}$ and an RNAP occupies the first $L_p$ bps of the gene,

$$U_{1:L_p}^{ON} \xrightarrow{k_{INI}} O_{1:L_p} \;.$$

In the next step an active RNAP, occupying nucleotide n to n+Lp-1, transcribes by taking one step forward with a rate $k_{EL}$ if the base $n+L_p$ is available, leaving a free base behind at location n,

$$O_{n:n+L_p-1} + U_{n+L_p} \xrightarrow{k_{EL}} O_{n+1:n+L_p} + U_n \;.$$

RNAP in active state ($O$) go to a paused state ($O^-$) with rate $k_{P+}$ and cease forward motion. A paused RNAP becomes active with a rate $k_{P-}$. The two reactions are given by the equations,

$$O_{n:n+L_p-1} \xrightarrow{k_{P+}} O_{n:n+L_p-1}^{-} \;,$$
$$O_{n:n+L_p-1}^{-} \xrightarrow{k_{P-}} O_{n:n+L_p-1} \;.$$

When an RNAP transcribes the last base of the gene it falls off leaving $L_p$ free bases and is modeled by the following reaction

$$O_{L-L_p+1:L} \xrightarrow{k_{EL}} U_{L-L_p+1:L} \;.$$

**Robustness of the Model**

In order to check the robustness of the results mentioned in the main text, we varied the key parameters of the model (hopping rate, gene length etc.) and found that the results hold for a wide range of parameters as shown in the following figures.

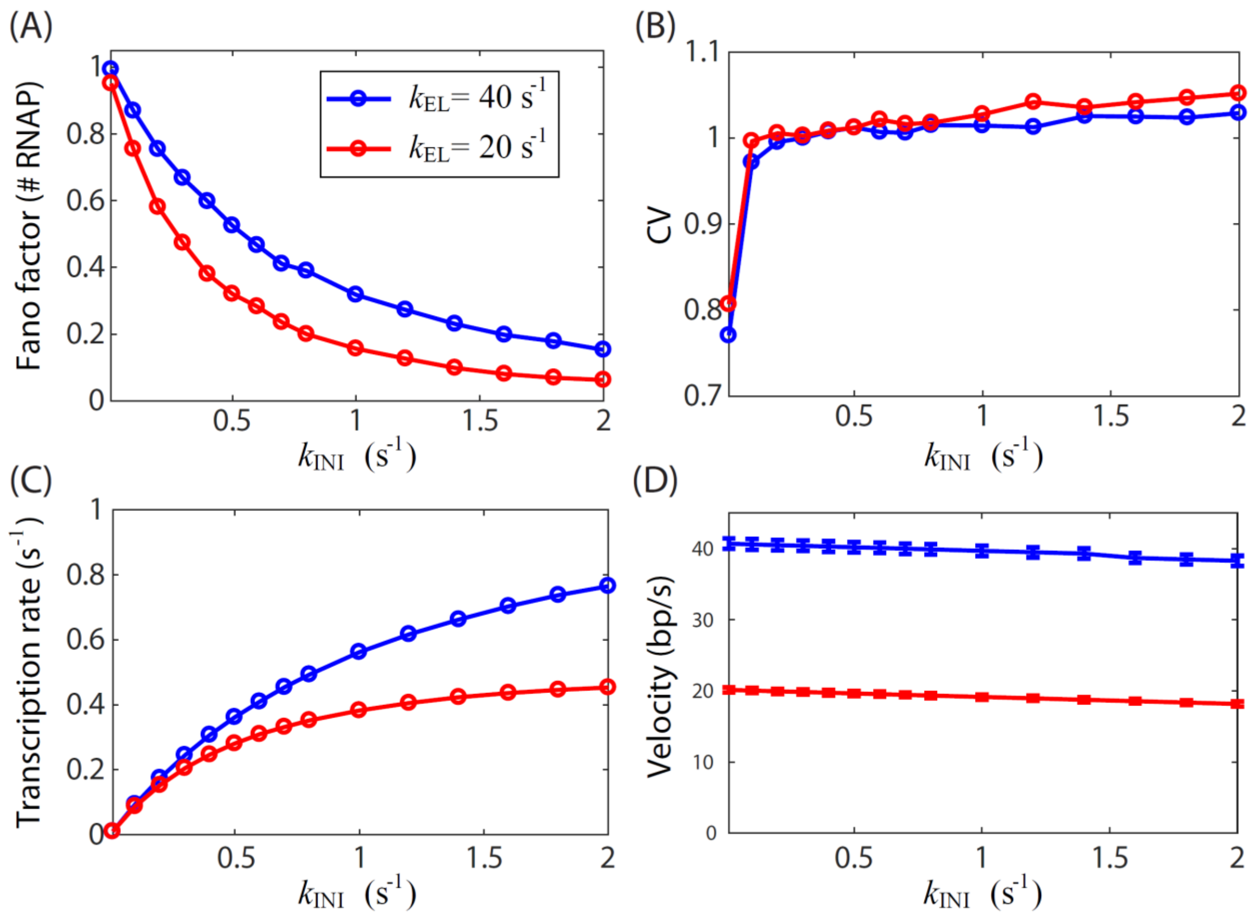


**Figure S1**

Noise profile in simple elongation model. **(A)** Fano factor of the RNAP number distribution and (**B)** CV of the inter-polymerase distance distribution along the gene, as a function of the initiation rate of the gene being transcribed for hopping rates, 20 s$^{-1}$ (red) and 40 s$^{-1}$ (blue). **(C)** The transcription rate and **(D)** average speed of polymerases along the gene of interest as a function of the initiation rate for hopping rates, 20 s$^{-1}$ (red) and 40 s$^{-1}$ (blue).

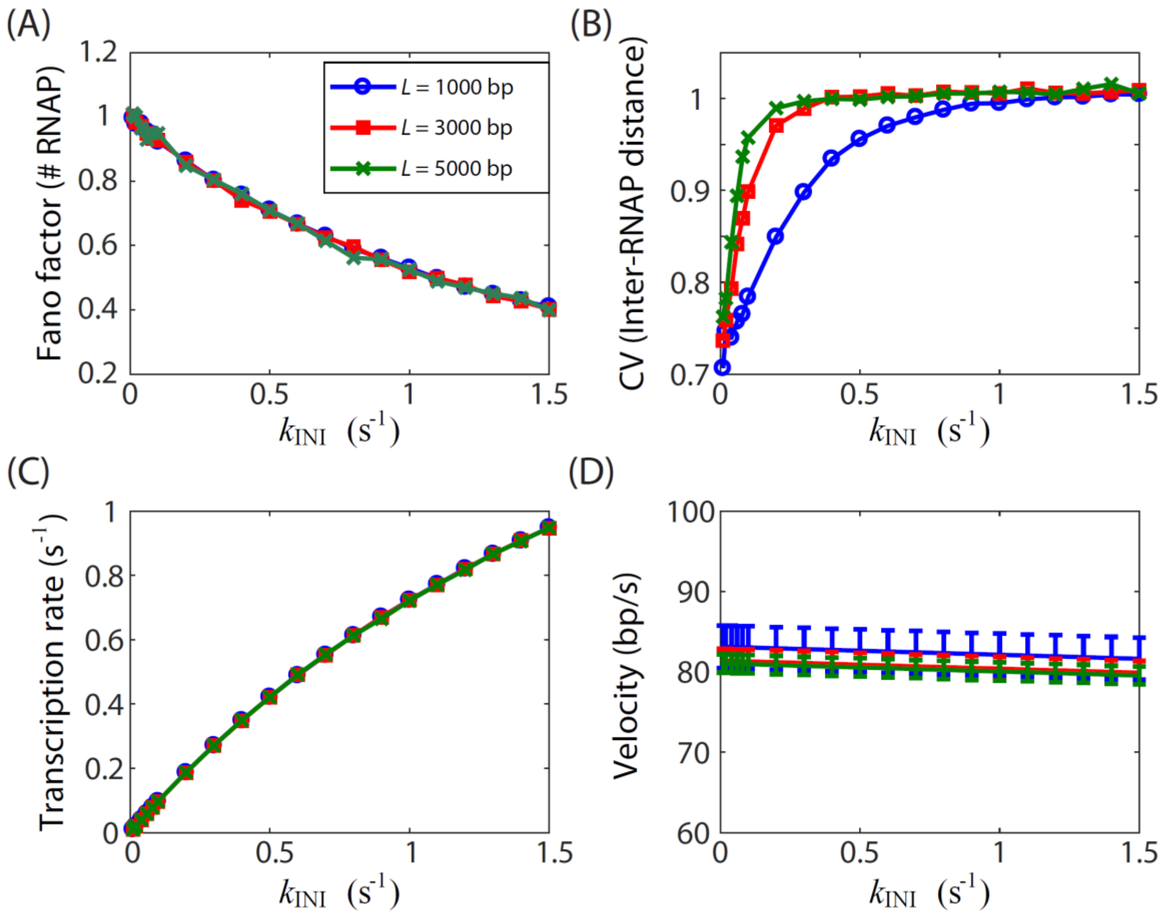


**Figure S2**

Noise profile in simple elongation model. **(A)** Fano factor of the RNAP number distribution and **(B)** CV of the inter-polymerase distance distribution along the gene, as a function of the initiation rate of the gene being transcribed for gene length 1000 bps (blue), 3000 bps (red), and 5000 bps (green). **(C, D)** The transcription rate and average speed of polymerases along the gene as a function of the initiation rate for gene length 3000 bps (blue), 3000 bps (red), and 5000 bps (green).

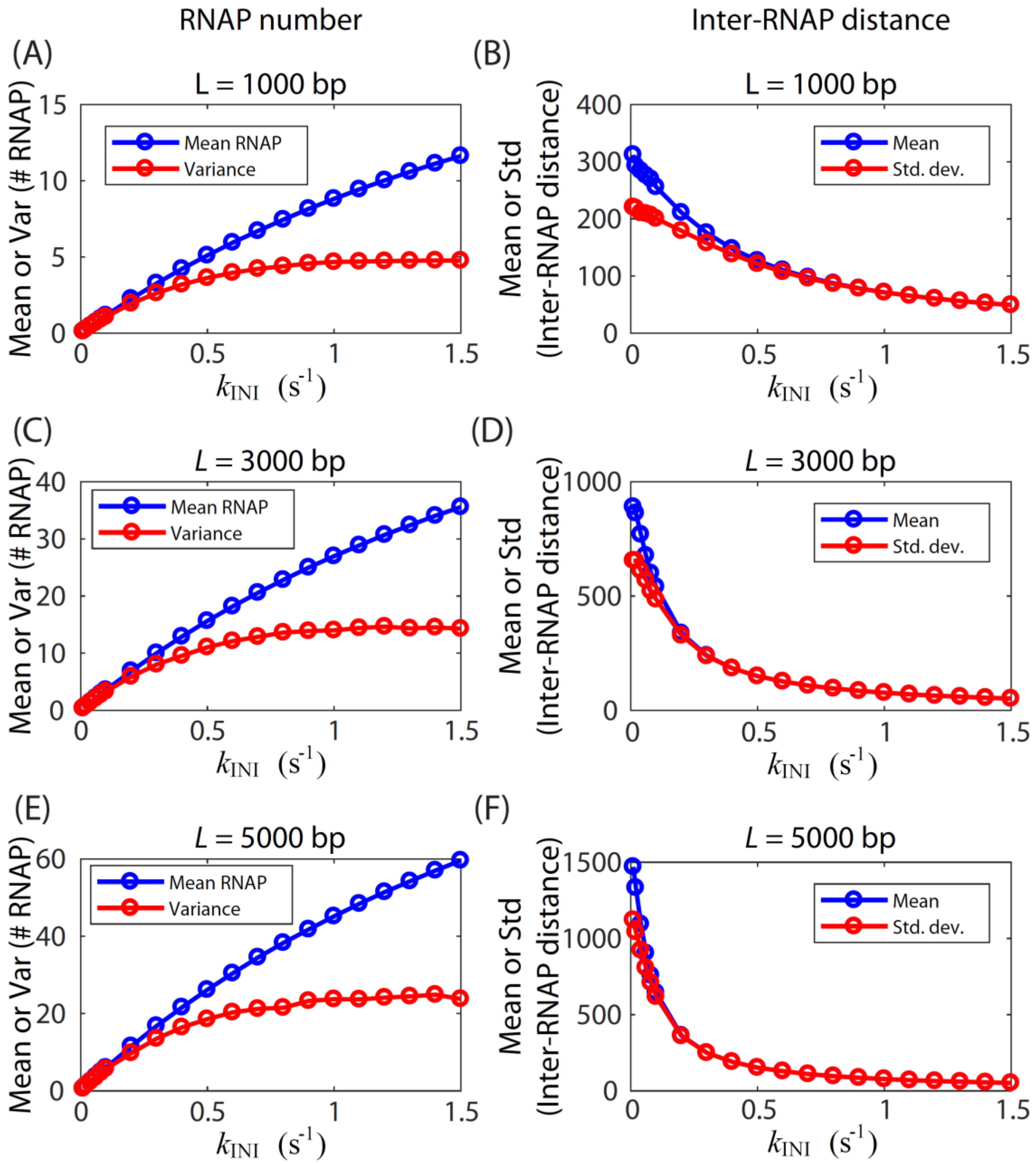


**Figure S3**

Noise profile in simple elongation model. **(A, C, E)** Mean (blue) and variance (red) of the RNAP number distribution on the gene, as a function of the initiation rate for gene lengths 1000 bps, 3000 bps, and 5000 bps. **(B,D,F)** Mean (blue) and standard deviation (red) of the inter-polymerase distance distribution along the gene for gene lengths 1000 bps, 3000 bps, and 5000 bps.

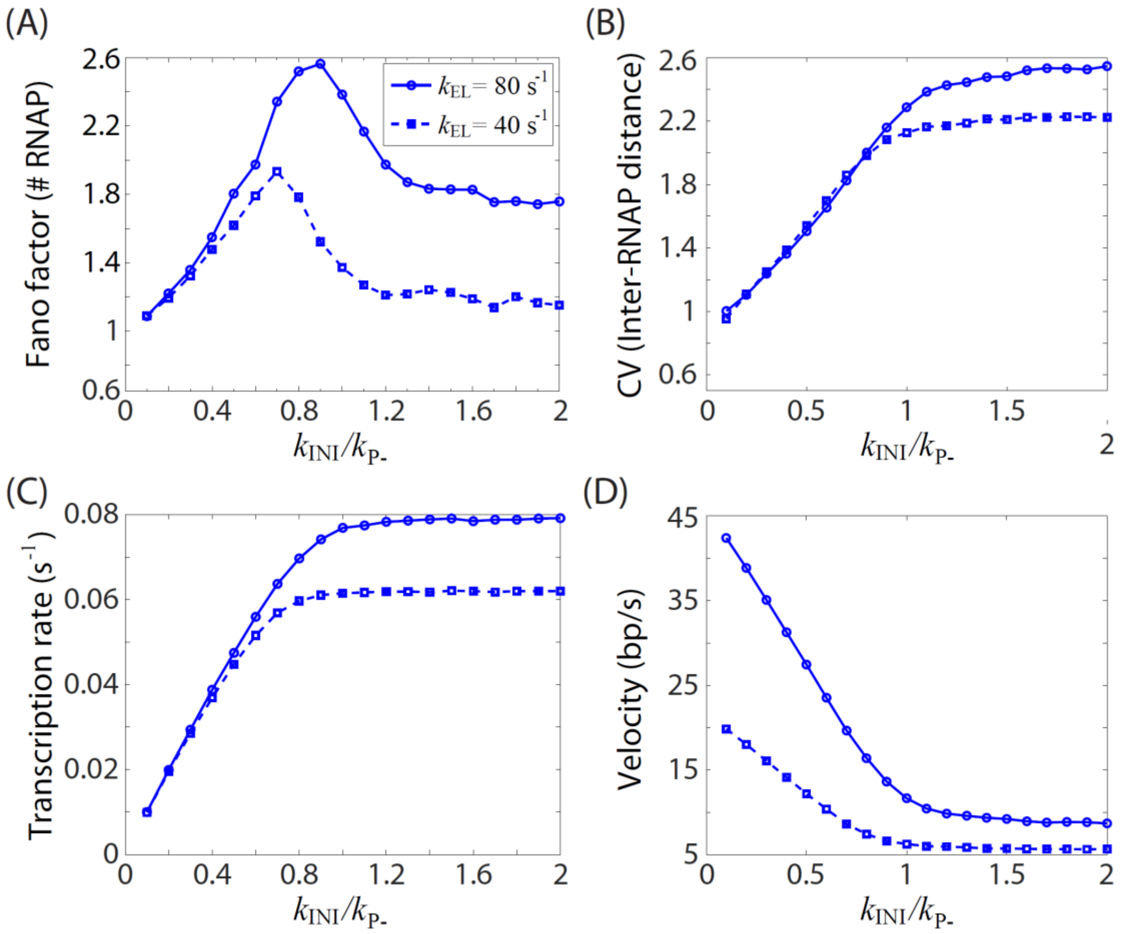


**Figure S4**

Noise profile in pausing model. (A,B) Fano factor of the nascent RNA number distribution and CV of the inter-polymerase distance distribution along the gene, as a function of the initiation rate of the gene being transcribed for hopping rates, 80 $s^{-1}$ (solid curve with circles) and 40 $s^{-1}$ (dashed curve with squares). (C, D) The transcription rate and average speed of polymerases along the gene of interest as a function of the initiation rate for hopping rates 80 $s^{-1}$ (solid curve with circles) and 40 $s^{-1}$ (dashed curve with squares).

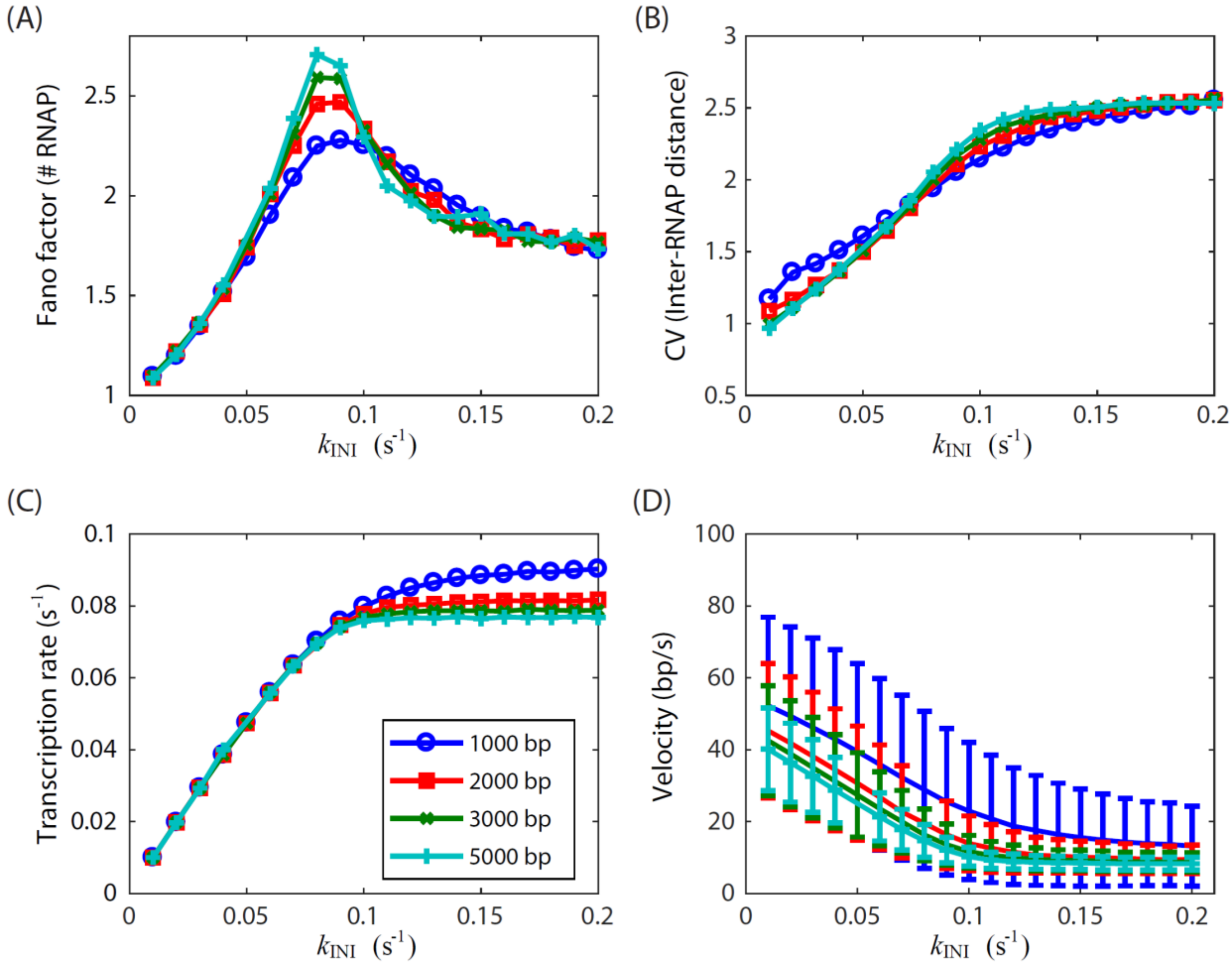


**Figure S5**

Noise profile in pausing model. (A,B) Fano factor of the RNAP number distribution and CV of the inter-polymerase distance distribution along the gene, as a function of the initiation of the gene being transcribed for gene lengths 1000 bps (blue), 2000 bps (red), 3000 bps (green), and 5000 bps (cyan). (C, D) The transcription rate and average speed of polymerases along the gene of interest as a function of the initiation rate for elongation rates.

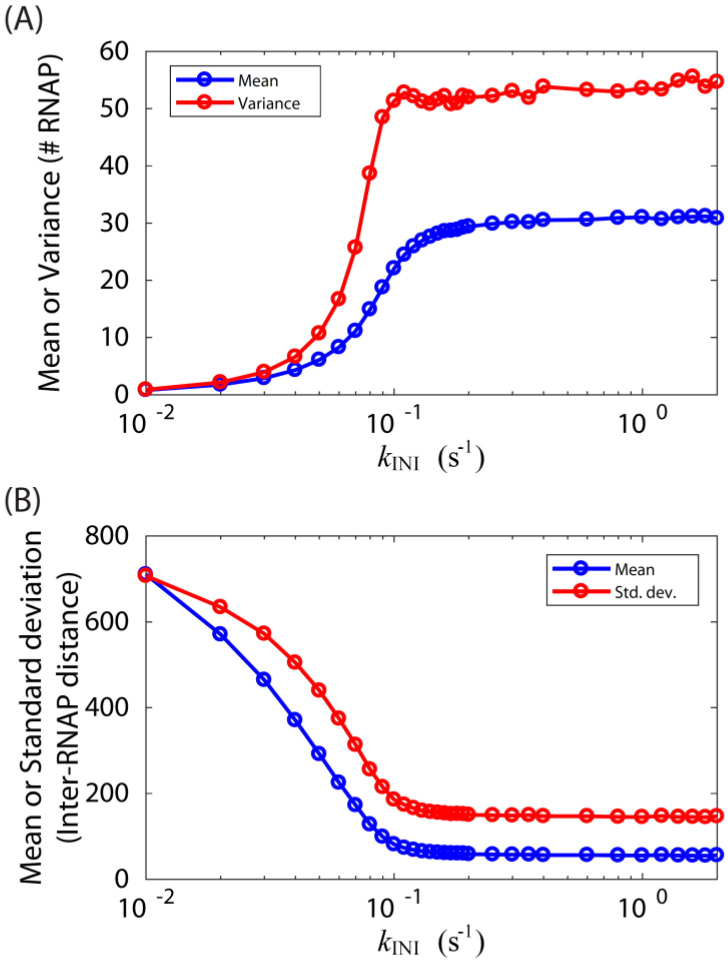


**Figure S6**

Noise profile in pausing model. (A) Mean (blue) and variance (red) of the RNAP number distribution on the gene, as a function of the initiation rate for gene length, $L$ = 3000 bps, $k_{P+}$ = 0.1 $s^{-1}$(red) and $k_{P-}$ = 0.1 $s^{-1}$(blue). (B) Mean (blue) and standard deviation (red) of the inter-polymerase distance distribution along the gene for the same parameters mentioned above.

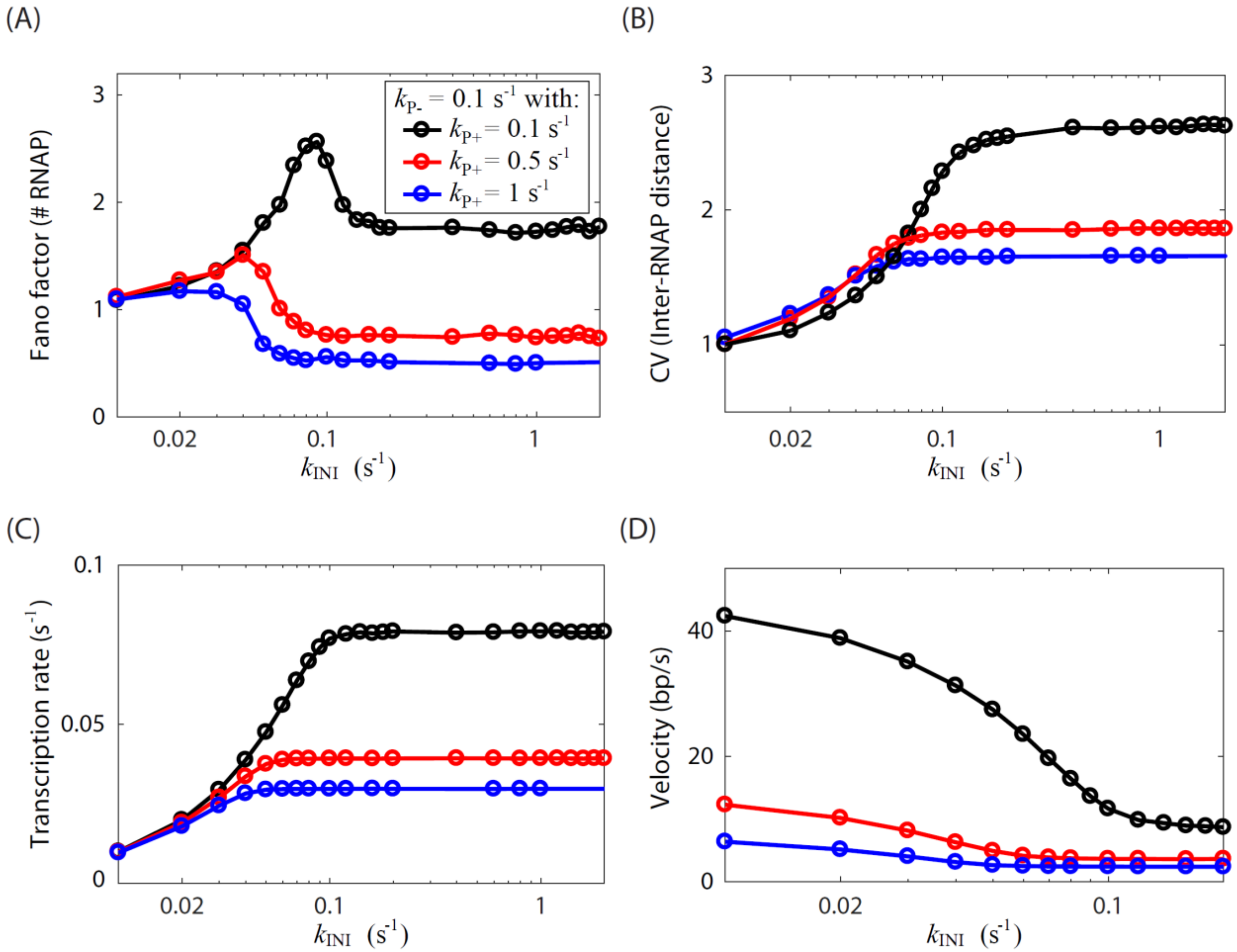


**Figure S7**

Noise profile in pausing model. (A,B) Fano factor of the RNAP number distribution and CV of the inter-polymerase distance distribution along the gene, as a function of the initiation of the gene being transcribed for $k_{P-}$ = 0.1 $s^{-1}$ and different $k_{P+}$ (0.1 $s^{-1}$, 0.5 $s^{-1}$, and 1 $s^{-1}$, black, red, and blue respectively). (C, D) The transcription rate and average speed of polymerases along the gene of interest as a function of the initiation rate for elongation rates.

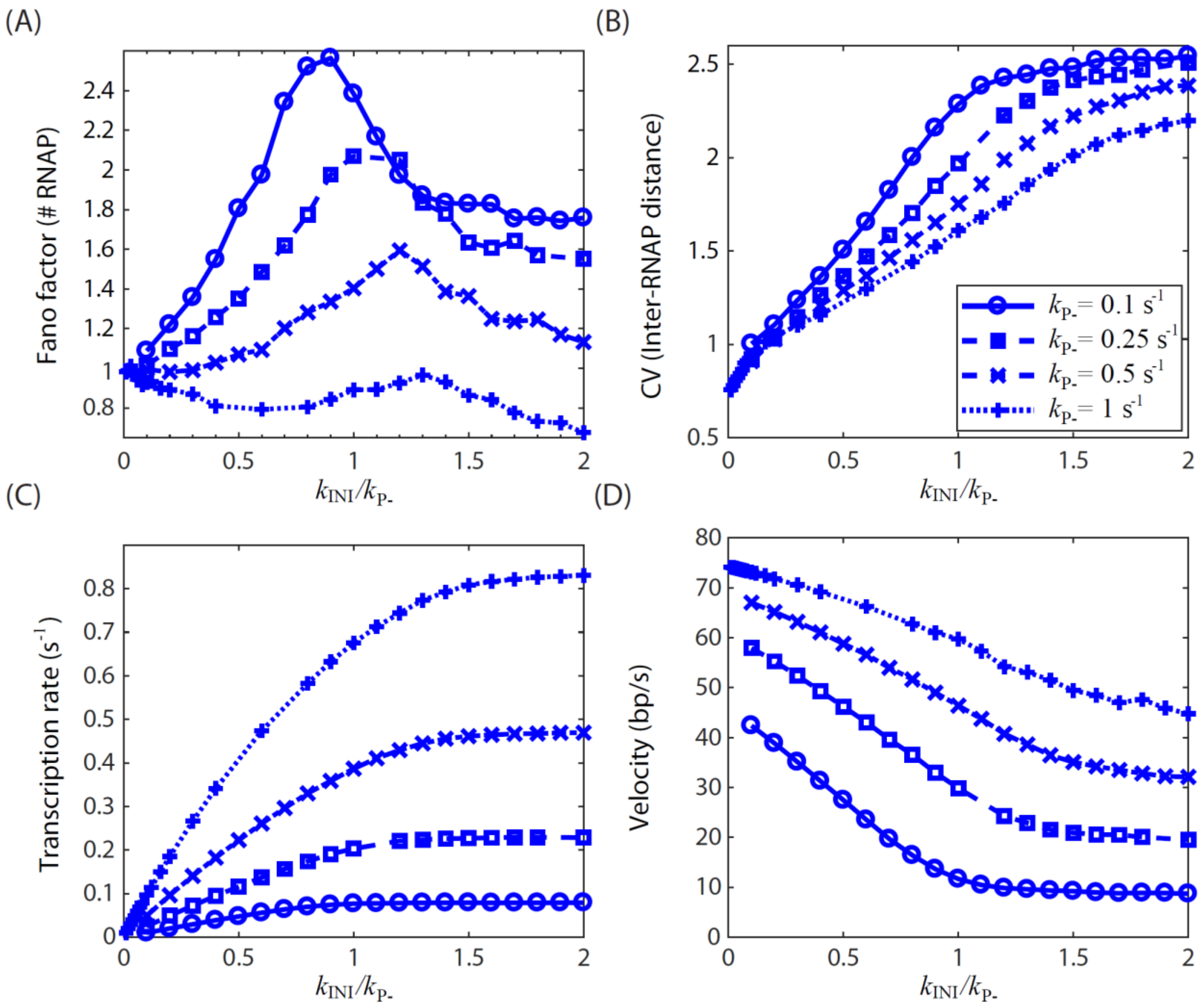


**Figure S8**

Noise profile in pausing model. (A,B) Fano factor of the RNAP number distribution and CV of the inter-polymerase distance distribution along the gene, as a function of the initiation over unpausing rate for $k_{P+}$ = 0.1 s$^{-1}$ and different $k_{P-}$ (0.1 s$^{-1}$, 0.25 s$^{-1}$, 0.5 s$^{-1}$, and 1 s$^{-1}$). (C, D) The transcription rate and average speed of polymerases along the gene of interest as a function of the initiation rate for elongation rates.

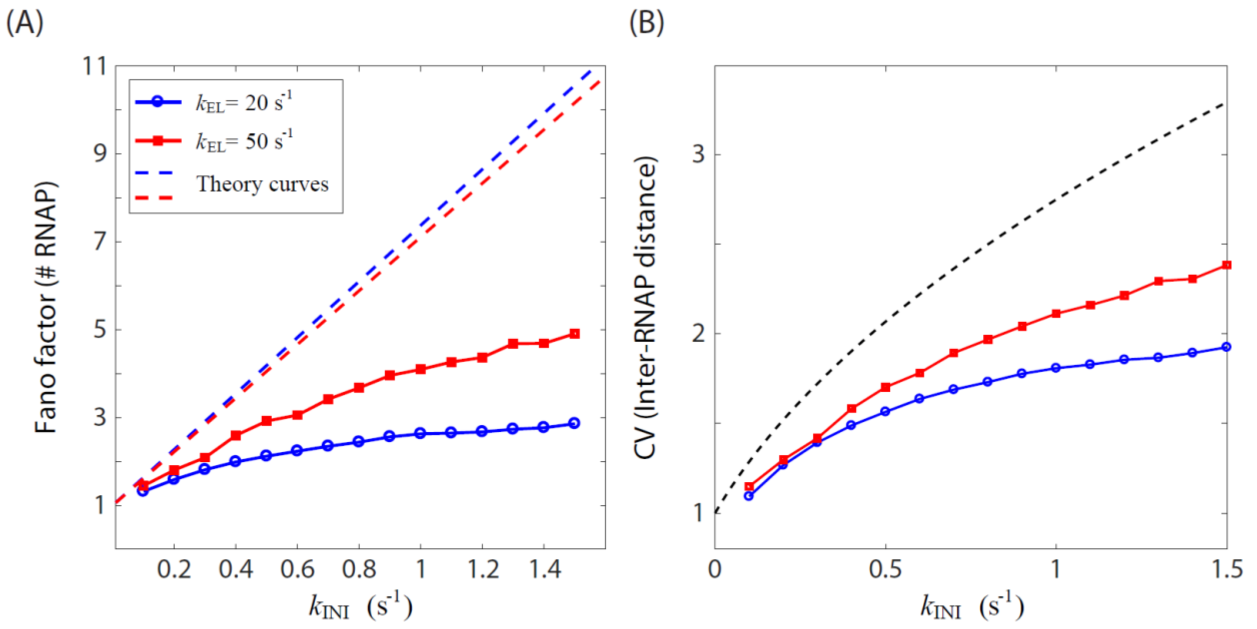


**Figure S9**

Noise profile in bursty initiation model followed by simple elongation. (A) Fano factor of the RNAP number distribution for hopping rates 20 s$^{-1}$ (blue) and 50 s$^{-1}$ (red). The dashed line represents the theoretical predictions from reference [1,3]. (B) CV of the inter-polymerase distance distribution along the gene, as a function of the initiation rate.

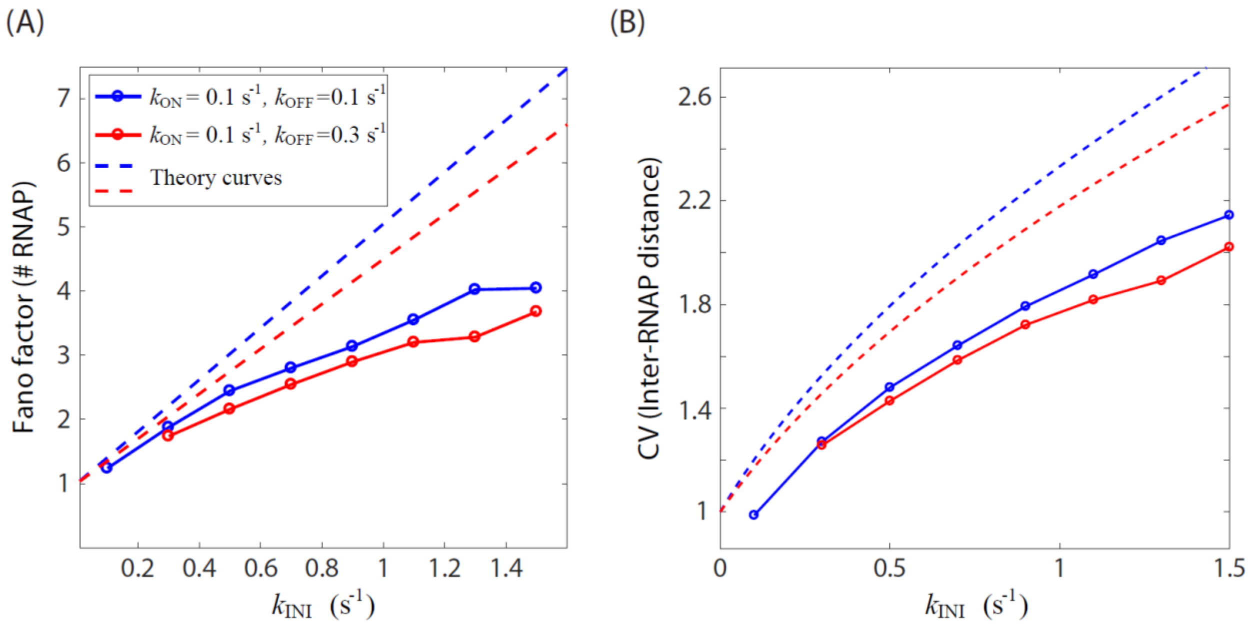


**Figure S10**

Noise profile in bursty initiation model followed by simple elongation. (A) Fano factor of the nascent RNA number distribution for $k_{ON}$ = 0.1 s$^{-1}$, $k_{OFF}$ = 0.1 s$^{-1}$ (blue) and $k_{ON}$ = 0.1 s$^{-1}$, $k_{OFF}$ = 0.3 s$^{-1}$ (red). The dashed line represents theoretical prediction from reference [1,3]. (B) CV of the inter-polymerase distance distribution along the gene, as a function of the initiation rate.

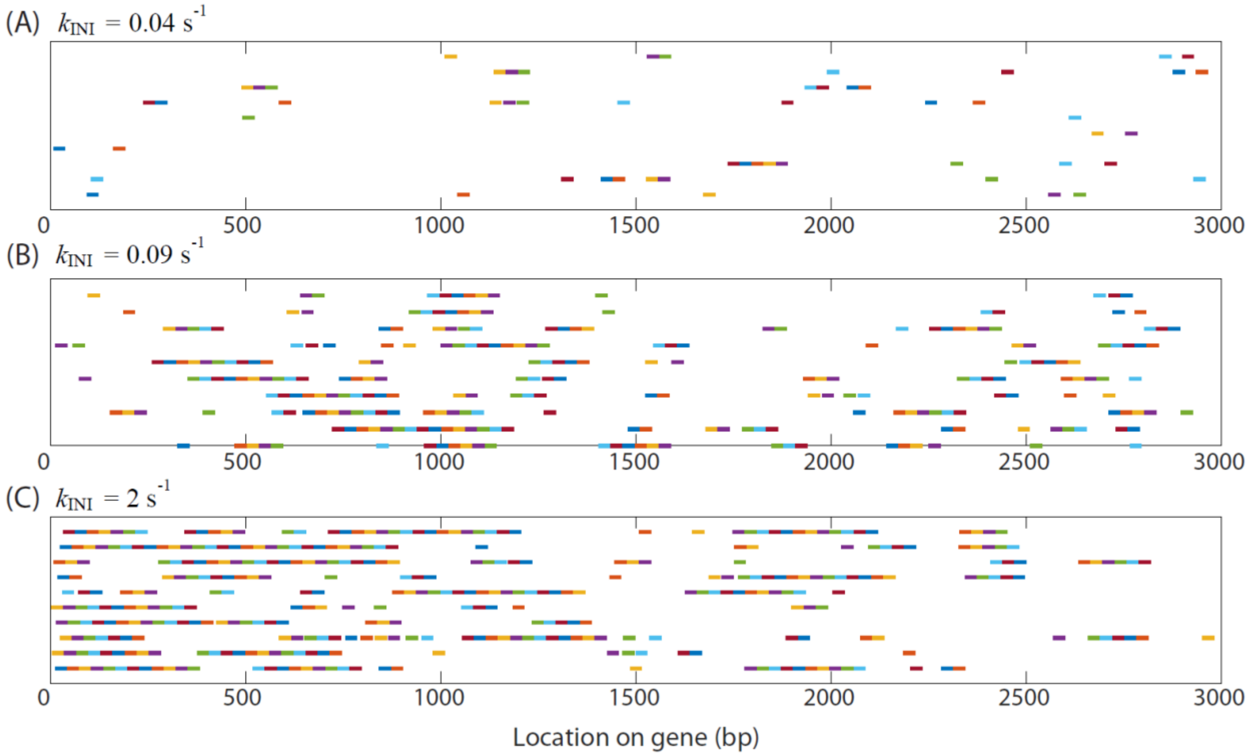


**Figure S11**
Snapshots from simulations showing bunches of RNAP molecules transcribing on a gene for the pausing model ($L$ = 3000 bp, $k_{EL}$ = 80 bp/s, $k_{P+}$ = 0.1 s$^{-1}$, $k_{P-}$ = 0.1 s$^{-1}$) for (A) $k_{INI}$ = 0.04 s$^{-1}$ , (B) 0.09 s$^{-1}$ , and (C) 2 s$^{-1}$. Each horizontal direction is a snapshot from a single simulation. RNAPs are denoted with different colors for distinction.

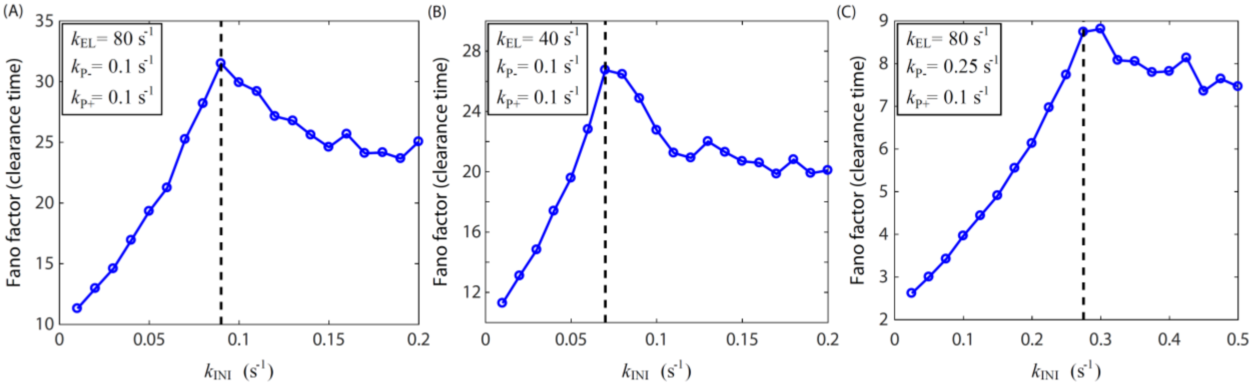


**Figure S12**
Fano factor for the time it takes for a polymerase molecule to traverse the gene for the Pausing model. The vertical dashed line indicates the initiation rates ($k_{INI}$) where the Fano factor for the RNAP number is maximum.

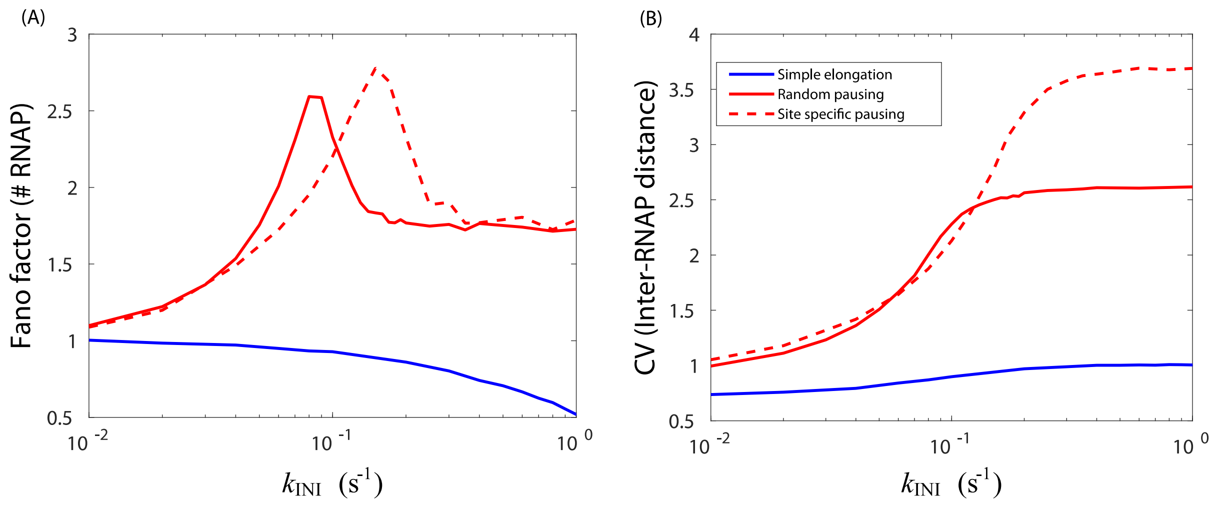


**Figure S13**

Site specific pausing versus pausing at every site for a constitutive promoter. **(A)** Fano factor of the RNAP number distribution and (**B)** CV of the inter-polymerase distance distribution along the gene, as a function of the initiation rate of the gene being transcribed for pausing at every site (solid red), site specific pausing (dashed red), and simple elongation (blue). For site specific pausing the RNAP molecules pauses at every 100 bps along the gene with pausing rate of $k_{P+}$ = 10/s which is 100 times larger than the pausing rate, $k_{P+}$ = 0.1/s, for random pausing at every site. The other parameters used are gene length $L$=3000 bps, hopping rate of 80 $s^{-1}$, and unpausing rate of $k_{P-}$ = 0.1/s.

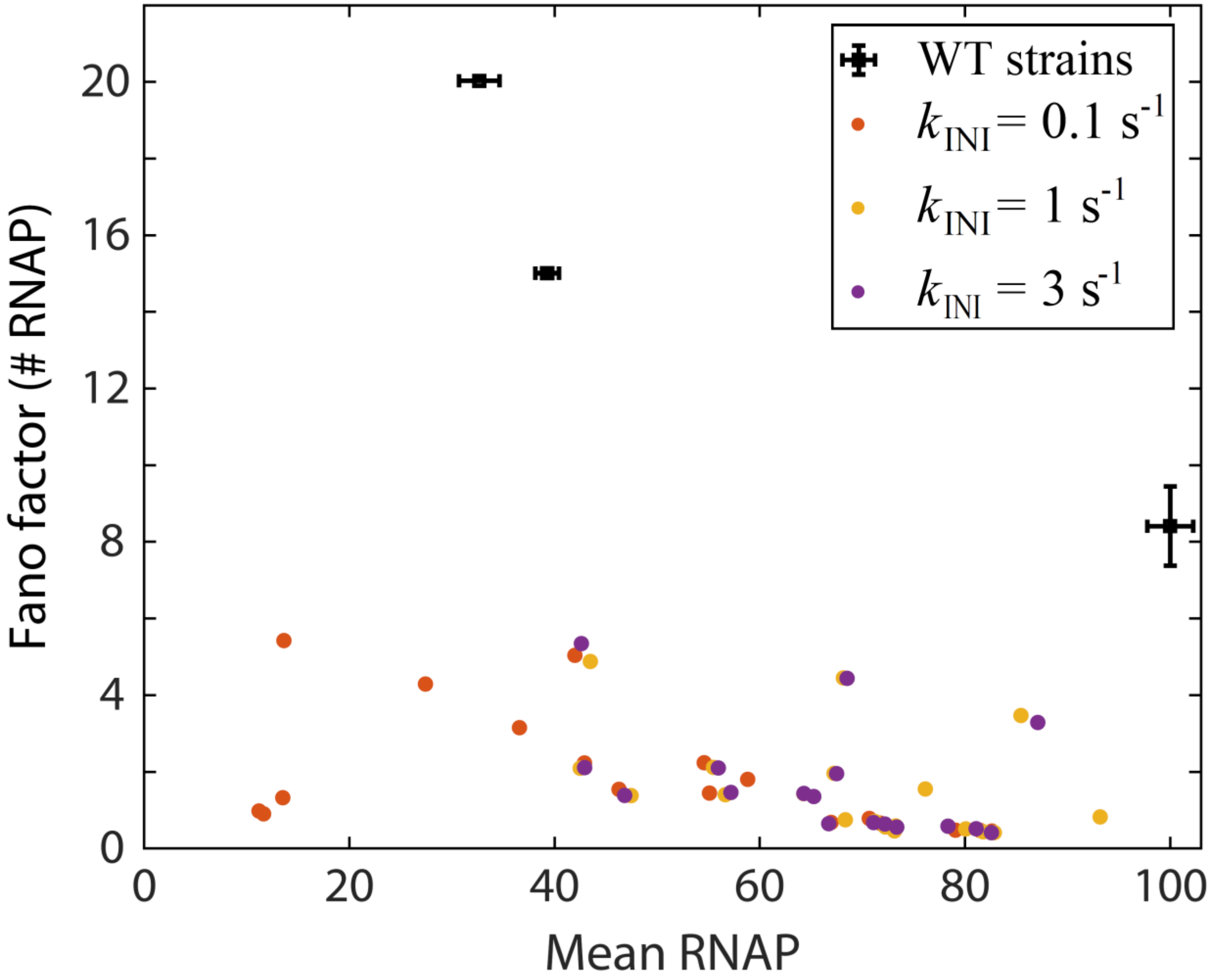


**Figure S14**

Fano factor of the RNAP number distribution in pausing model as a function of mean RNAP number by varying the initiation rate ($k_{INI}$ = 0.1, 1, and 3 s$^{-1}$), pausing rate ($k_{P+}$ = 0.01, 0.05, 0.1, 0.5, and 1 s$^{-1}$), and unpausing rate ($k_{P-}$ = 0.01, 0.05, 0.1, and 1 s$^{-1}$). The Fano factor for wild type (WT) strains from experimental studies [4–6] are shown in black square with error bars.

**References**


1. Choubey S, Kondev J, Sanchez A. Deciphering Transcriptional Dynamics In Vivo by Counting Nascent RNA Molecules. PLOS Comput Biol. 2015;11: e1004345. doi:10.1371/journal.pcbi.1004345
2. Choubey S. Nascent RNA kinetics: Transient and steady state behavior of models of transcription. Phys Rev E. 2018;97: 022402. doi:10.1103/PhysRevE.97.022402
3. Choubey S, Kondev J, Sanchez A. Distribution of Initiation Times Reveals Mechanisms of Transcriptional Regulation in Single Cells. Biophys J. 2018;114: 2072–2082. doi:10.1016/j.bpj.2018.03.031
4. Claypool JA, French SL, Johzuka K, Eliason K, Vu L, Dodd JA, et al. Tor Pathway Regulates Rrn3p-dependent Recruitment of Yeast RNA Polymerase I to the Promoter but Does Not Participate in Alteration of the Number of Active Genes. Mol Biol Cell. 2004;15: 946–956. doi:10.1091/mbc.E03-08-0594
5. Tongaonkar P, French SL, Oakes ML, Vu L, Schneider DA, Beyer AL, et al. Histones are required for transcription of yeast rRNA genes by RNA polymerase I. Proc Natl Acad Sci. 2005;102: 10129–10134. doi:10.1073/pnas.0504563102
6. Schneider DA. Quantitative analysis of transcription elongation by RNA polymerase I in vitro. Methods Mol Biol Clifton NJ. 2012;809: 579–591. doi:10.1007/978-1-61779-376-9_37
